# Attention differentially modulates the amplitude of resonance frequencies in the visual cortex

**DOI:** 10.1101/518779

**Authors:** Rasa Gulbinaite, Diane H. M. Roozendaal, Rufin VanRullen

**Affiliations:** Lyon Neuroscience Research Center (CRNL), Brain Dynamics and Cognition Team, INSERM U1028, CNRS UMR5292, Université Claude Bernard Lyon 1, UdL, Lyon, France; Department of Psychology, Department of Brain and Cognition, University of Amsterdam, Netherlands; Centre National de la Recherche Scientifique, UMR 5549, Faculté de Médecine Purpan, Toulouse, France; Université de Toulouse, Centre de Recherche Cerveau et Cognition, Université Paul Sabatier, Toulouse, France

## Abstract

Rhythmic visual stimuli (flicker) elicit rhythmic brain responses at the frequency of the stimulus, and attention generally enhances these oscillatory brain responses (steady state visual evoked potentials, SSVEPs). Although SSVEP responses have been tested for flicker frequencies up to 100 Hz [Herrmann, 2001], effects of attention on SSVEP amplitude have only been reported for lower frequencies (up to ~30 Hz), with no systematic comparison across a wide, finely sampled frequency range. Does attention modulate SSVEP amplitude at higher flicker frequencies (gamma band, 30-80 Hz), and is attentional modulation constant across frequencies? By isolating SSVEP responses from the broadband EEG signal using a multivariate spatiotemporal source separation method, we demonstrate that flicker in the alpha and gamma bands elicit strongest and maximally phase stable brain responses (resonance), on which the effect of attention is opposite: positive for gamma and negative for alpha. Finding subject-specific gamma resonance frequency and a positive attentional modulation of gamma-band SSVEPs points to the untapped potential of flicker as a non-invasive tool for studying the causal effects of interactions between visual gamma-band rhythmic stimuli and endogenous gamma oscillations on perception and attention.

## INTRODUCTION

More than twenty years ago, Morgan and colleagues (1996) elegantly demonstrated that attended vs. ignored flickering stimuli elicit stronger periodic cortical activity in EEG (steady-state visual evoked potentials, SSVEPs): A finding that allowed measuring attention continuously over space and time without the need for explicit responses from participants. Although SSVEP responses were tested for flicker frequencies up to 100 Hz (Herrmann, 2001), the effects of attention on SSVEP amplitude only have been reported for lower frequencies (up to ~30 Hz, Kashiwase et al., 2012; Kus et al., 2013), with no systematic comparison across flicker frequencies. Does attention modulate SSVEP amplitude to flicker frequencies in the range of endogenous gamma-band oscillations (30-80 Hz), which are implicated in attention but difficult to study in human EEG (Fries et al., 2008; Hipp and Siegel, 2013; Hoogenboom et al., 2006)? Is the effect of attention on SSVEP amplitude constant or variable across a wide-range of flicker frequencies (3-80 Hz)? How do flicker responses interact with endogenous rhythms implicated in attentional processes (Frey et al., 2015; Green et al., 2017)?

Attending to flickering stimuli generally enhances SSVEP amplitude (Kashiwase et al., 2012; Kus et al., 2013; Morgan et al., 1996; Muller et al., 1998b). However, attention effects can be reversed for flicker frequencies in the low alpha band (7-10 Hz): The SSVEP amplitude was decreased for attended as compared to ignored stimuli (7.41 and 8.6 Hz in Chen et al., 2003; 9.2 Hz in Ding et al., 2006). Although Ding and colleagues deemed unlikely that endogenous alpha oscillations could contribute to the reversed attention effect of alpha-band SSVEPs (Ding et al., 2006), it is plausible to assume that at the electrode level SSVEPs represent a mixture of responses to flicker and alpha-band frequency components of endogenous EEG (Onton and Makeig, 2006). Attention-related modulations of endogenous alpha-band power are well-documented (for reviews, see Foxe and Snyder, 2011; Frey et al., 2015; Klimesch et al., 2007), and consistently show decreases in alpha-band power related to visual stimulation and increased attention. Despite accumulating evidence for behaviorally-relevant alpha-band flicker effects on attention (Gulbinaite et al., 2017; Kizuk and Mathewson, 2017; Shapiro et al., 2017), with the strongest effects when flicker rate matches individual alpha peak frequency (de Graaf et al., 2013; Gulbinaite et al., 2017), electrophysiological evidence for entrainment of endogenous oscillations using rhythmic sensory stimulation is still an active area of research (Capilla et al., 2011; Norcia et al., 2015; Thut et al., 2011; Zoefel et al., 2018). Characterization of endogenous oscillations and simultaneously co-occurring SSVEPs poses a non-trivial signal processing problem because of the overlapping cortical foci of the two signals (Spaak et al., 2014). To isolate frequency-specific SSVEPs from the broadband EEG signal, we used a guided multivariate spatiotemporal source separation (Cohen and Gulbinaite, 2017).

In this study, using a covert spatial attention task and a custom-built LED-based hardware to ensure precision of stimulus timing, we sought to map out attentional modulation of SSVEPs using a wide, finely sampled range of tagging frequencies (3-80 Hz). We focused on the following specific aims. First, we aimed to determine whether attention modulates SSVEP amplitude at higher flicker frequencies (above 30 Hz), and to test whether attentional modulation of SSVEPs is constant across frequencies. Second, we aimed to characterize the effects of attention simultaneously on endogenous alpha oscillations vs. alpha-band SSVEPs. Third, we investigated attentional modulation of brain responses at 2*f*, twice the stimulus frequency (second harmonic), which can also be modulated by attention, and sometimes show stronger attentional modulations as compared to the stimulus frequency (Kim et al., 2007; Kim et al., 2011; Vissers et al., 2017).

We also tested whether stability of SSVEP phase over time – a feature commonly attributed to SSVEPs (Norcia et al., 2015; Vialatte et al., 2010) – is a valid assumption, and whether phase stability of SSVEPs is frequency dependent. Classical SSVEP analysis is focused on amplitude and phase at the stimulation frequency, which are determined by applying the Fourier transform to the trial-average waveforms, or, in case of single-trial analyses, to the running average waveforms (Nunez and Srinivasan, 2006; Wieser et al., 2016). Such approaches extract only the phase-locked part of the SSVEP signal, and remove non-stationary features of SSVEPs captured by the side-bands in the frequency spectrum. To capture fluctuations in the phase of SSVEP response within each trial without the averaging step, we used the first temporal derivative of the phase time series (Cohen, 2014).

We found statistically robust responses to flicker up to 80 Hz, with participant-specific resonance response peaks to flicker in the alpha (~10 Hz) and gamma (~47 Hz) bands. The sign of attentional modulation of SSVEP amplitude was frequency dependent: Positive for theta- (3-7 Hz) and gamma-band (30-80 Hz) flicker frequencies, but negative for alpha (8-13 Hz). By characterizing the effects of attention simultaneously on endogenous alpha oscillations and alpha-band SSVEPs, we demonstrate that attention-related changes in endogenous alpha is the most plausible cause for amplitude reversal of alpha-band SSVEPs. Furthermore, we provide evidence that SSVEP frequency varies over time, and most phase-stable SSVEPs are generated in response to alpha- and gamma-band flicker.

## METHODS

### Participants

Twenty participants took part in this experiment in exchange for a monetary compensation (€10/hour). The required sample was determined based on our previous studies using a frequency tagging approach to study attention (Gulbinaite et al., 2014; Gulbinaite et al., 2017), and those used in previous reports using similar task designs (Keitel et al., 2013; Toffanin et al., 2009). All had normal or corrected-to-normal vision, were screened for photosensitivity and family history of epilepsy or migraine. The experiment was conducted in accordance with the Declaration of Helsinki and approved by the local ethics committee. Written informed consent was obtained from all participants before the experiment. The data from one participant were excluded from the analyses due to excessive movement artifacts in the EEG (>50% of the trials had to be discarded). Thus, the final sample was 19 participants (5 female; mean age of 27.35, SD = 3.60).

### Experimental setup

We used a custom-built setup with arrays of light-emitting diodes (LED) to deliver visual stimulation (Figure 1A). The setup consisted of two main parts: (1) single white fixation LED positioned centrally at the level of the eyes (luminous intensity 11000 mcd, color temperature 3500 K, Model NSPL500DS, Nichia Corporation); (2) two arrays of 12 LEDs, each containing 4 blue LEDs (luminous intensity 11000 mcd, Model C503B-BAS, Cree Inc.) surrounded by 8 white LEDs (luminous intensity 6900 mcd, color temperature 9000K, Model C513A-WSN, Cree Inc.). For diffusion of light, LED arrays and fixation LED were placed in cardboard boxes, with the apertures covered by tracing paper (6 cm away from LED arrays). The apertures for fixation and stimulus circles subtended 0.87° and 5.2° of visual angle, respectively. The stimulus LED array was placed 5.2° below the horizontal meridian at an eccentricity of 4.3°. The luminance of white and blue LEDs was linearized using a quadratic function, such that a linear increase in voltage corresponded to a linear increase in luminance (measured in cd/m^2^).

**Figure 1.**
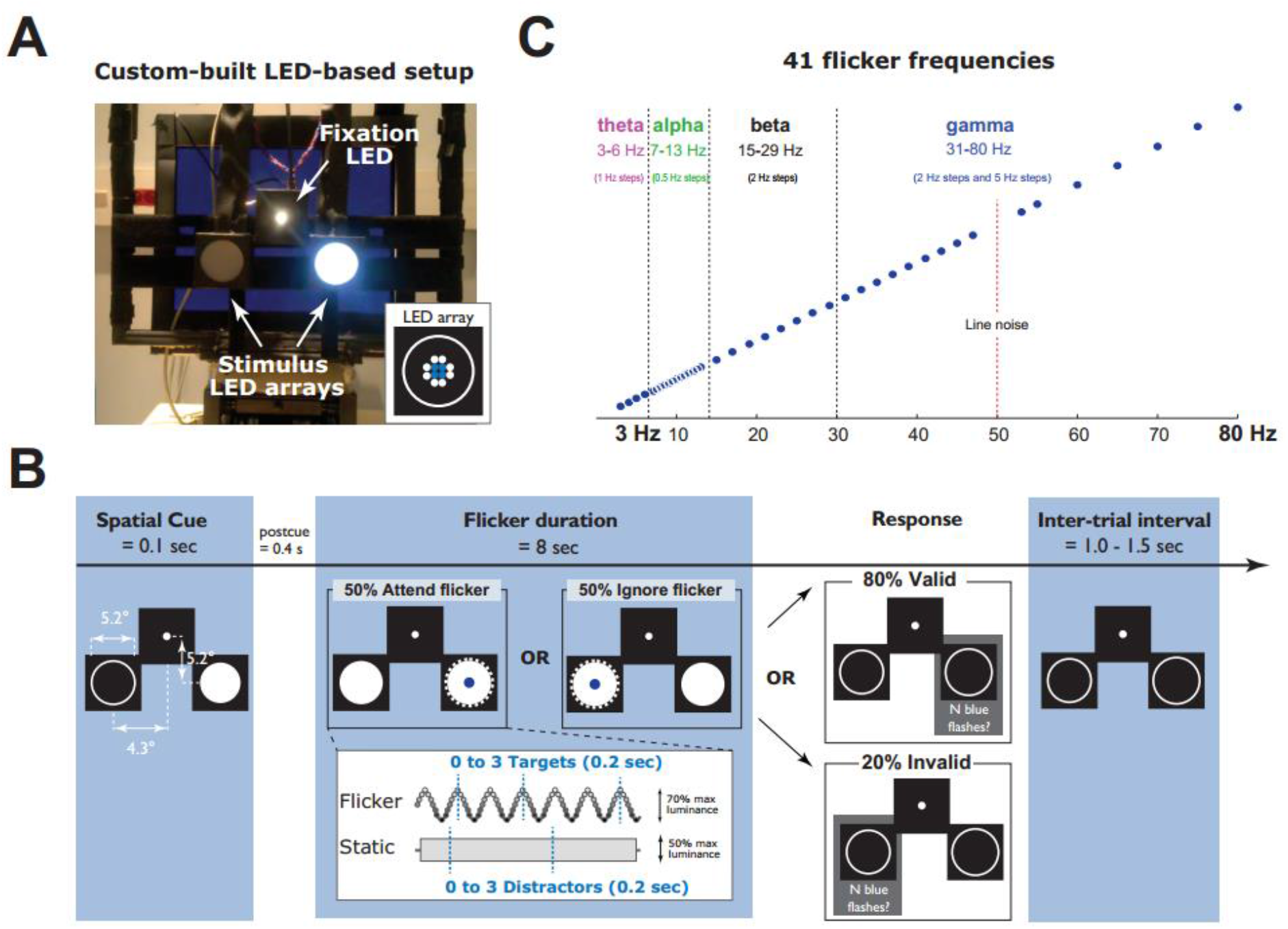
Visual stimulation setup and trial structure. (A) LED-based setup with fixation LED and stimulus LED arrays (spatial configuration of white and blue LEDs is represented in the inset) were placed in front of the monitor screen. (B) Each trial started with a cue (80% validity), which indicated the to-be-attended location for that trial (left or right). Following the post-cue interval, for a duration of 8000 ms the luminance of white LEDs in one of the locations was modulated by a sine-wave at a constant frequency (randomly selected from 41 possible frequencies ranging from 3 to 80 Hz), the luminance of white LEDs on the other side was kept constant. This corresponded to “attend flicker” and “ignore flicker” conditions. Zero to three blue flashes could appear on both sides, and subjects were instructed to count the number of blue flashes at the cued location. Following 8000 ms, a grey square was presented at one of the locations, indicating to report the number of blue flashes presented on that side. Following the response, the next trial started after a variable delay (1000-1500 ms). (C) Flicker frequency distribution. In total, 41 flicker frequencies ranging from 3 to 80 Hz were used: 4 frequencies in the theta band (3-6 Hz, 1 Hz step), 13 frequencies in the alpha band (7-13 Hz, 0.5 Hz step), beta band (15-29 Hz, 2 Hz step), and gamma band (31-80 Hz, steps of 2 Hz and 5 Hz).

Flicker was implemented as a sine-wave modulation of the power supply to the LEDs, which was controlled using a microcontroller of the STM32F4-Discovery board by adjusting the width of PWM signal, and linearizing the current using a low-pass filter. MATLAB was used for communication with the STM32F4-Discovery board via a USB virtual com port protocol (programmed in C++). A circuit diagram of the sinewave generator can be found in Supplementary materials (Figure S1). Luminance of the white LEDs in the stimulus LED-array was kept constant throughout the experiment, whereas the luminance of blue LEDs was adjusted using a staircase procedure and was on average 253 cd/m^2^ (see Stimuli and Procedure for further details). Participants were seated in a dark room. The viewing distance was constrained by the chin rest positioned 66 cm away from the custom-built LED setup. The setup was attached and placed in front of a 17-inch CRT monitor (1280 x 1024 pixels; 85Hz refresh rate).

### Stimuli and procedure

A covert visuospatial attention task was used to manipulate spatial attention. Participants were instructed to focus on the fixation LED throughout the experiment, covertly shift their attention based on the cue location, and count the number of blue flashes (targets) in the cued hemifield, while ignoring appearance of the blue flashes (distractors) in the uncued hemifield. A spatial cue (80% validity, randomized across trials) indicated the to-be-attended hemifield (50% left, randomized across trials).

Each trial began with a presentation of the spatial cue: One of the white LED arrays was turned on for 100 ms, luminance level of 996 cd/m^2^. Following a 400 ms post-cue interval, one of the white LED arrays started flickering (sine-wave modulated luminance changes from 0 to 1970 cd/m^2^) at one of the 41 possible frequencies (ranging from 3 to 80 Hz), while the luminance of the other white LED array was kept constant (2530 cd/m^2^) for the duration of 8000 ms (Figure 1B). Thus, half of the trials were “attend flicker” and half were “ignore flicker”. At the end of each trial, a grey square appeared on the monitor screen behind the left or the right LED array box, indicating to report the number of blue flashes presented on that side by pressing the appropriate button on the numpad of the keyboard. Location of the grey box matched the location of the spatial cue on 80% trials (valid trials), on the remaining 20% of trials the grey box appeared in the uncued hemifield (invalid trials).

On each trial, 0-3 targets (blue-flashes in the cued hemifield) and 0-3 distractors (blue-flashes in the uncued hemifield) appeared at random times for the duration of 200 ms. The number of targets and distractors was randomly drawn from a gamma distribution (k = 7, and θ = 0.35). To achieve maximal unpredictability of the next stimulus onset, the timing of the blue flashes was determined using three truncated negative exponential distributions based on the number of blue flashes: (1) on single-flash trials, the blue flash occurred in the 1-8 sec interval relative to the flicker onset (λ = 3); (2) on two-flash trials, the 1^st^ flash occurred in the 1-3 sec interval relative to the flicker onset (λ = 2), interstimulus interval (ISI) between 1^st^ and 2^nd^ blue flash was minimum 1 second (λ = 1); (3) on three-flash trials, the 1^st^ flash occurred in the 1-3 sec interval relative to the flicker onset (λ = 2), the 2^nd^ flash occurred with the ISI of 1-6 sec relative to the 1^st^ flash (λ = 1), the 3^rd^ flash occurred with the ISI of 1 sec (λ = 1).

Performance accuracy was kept at 80% on valid trials by adjusting the luminance of the blue LEDs using a QUEST procedure with a sliding window of 40 trials, separately for “attend flicker” and “ignore flicker” (attend static white LEDs) trials. The luminance of the blue LEDs was also adjusted taking into account flicker frequency, with a frequency window for the QUEST procedure set to +-5 frequencies surrounding the frequency of interest. This was done to accommodate variability in perceived stimulus luminance as a function of the flicker frequency (Bertrand et al., 2018; Wu et al., 1996).

In total, we used 41 logarithmically spaced flicker frequencies that ranged from 3 to 80 Hz, with denser sampling in the alpha-band (7-14 Hz in steps of 0.5 Hz; Figure 1C). Each frequency was presented on 12 trials: Three times in four different conditions (attend right flicker, ignore right flicker, attend left flicker and ignore left flicker), counterbalanced and randomized across trials. The experiment consisted of 10 practice trials and 492 experimental trials divided into 12 blocks. Feedback on performance accuracy was provided after each trial during the practice session. In the experimental session, average response accuracy was provided after each block. Participants were encouraged to take self-paced breaks during the experiment and took mandatory breaks after each block. The experiment lasted ~2.5 h. Resting state eye-open and eyes-closed EEG recording (2 minutes each) was obtained prior to the beginning of the experiment.

Eye movements were recorded throughout the experiment using EyeLink 1000 Plus (version 5.04, SR Research, Ontario) with a sampling rate of 1000 Hz, and 9-point custom calibration. Gaze data were analyzed offline using EyeLink^®^ Data Viewer to identify the trials during which participants made saccades to one of the lateralized stimuli during the time window used for SSVEP analysis. No trials were rejected based on the inspection of eye movement data. Stimulus presentation, response registration, and interaction with EyeLink 1000 Plus were controlled by custom-written Matlab routines using Psychtoolbox (Brainard, 1997).

### EEG data acquisition and preprocessing

EEG data were recorded using a 64-channel ActiveTwo BioSemi system (for detailed description of the hardware, see www.biosemi.com) at 1024 Hz sampling rate. Offline, the data were downsampled to 512 Hz and re-referenced to the average activity of all electrodes. Continuous EEG data were high-pass filtered at 0.5 Hz. The data were then epoched (−1000 to 9000 ms relative to the cue onset) and baseline-corrected to the −200 – 0 ms time window. Upon visual inspection of the data, bad electrodes and trials containing excessive movement artifacts were rejected (short movement artifacts are not detrimental for SSVEP analyses, Norcia et al., 2015). Eyeblink-related artifacts were identified using independent component analysis ICA using the JADE algorithm (Delorme and Makeig, 2004). Independent components (ICs) representing eyeblinks were identified based on their frontal topography and relatively high power at the low frequencies (following criteria provided by Chaumon et al., 2015), and were removed only for analyses involving alpha-band power, and 3-6 Hz flicker conditions. Analyses involving spatiotemporal source-separation (described in the following section) did not require removing artifactual independent components and were done on the full-rank data.

### Construction of frequency-specific SSVEP spatiotemporal filters

Frequency-specific SSVEP responses were obtained using a spatiotemporal source separation method called rhythmic entrainment source separation (RESS; Cohen and Gulbinaite, 2017). The method exploits the temporal (flicker frequency) and spatial (topographical distribution) characteristics of SSVEPs to determine an optimal spatial filter (channel weights) that maximally separates frequency-specific SSVEPs (signal, **S**) from the brain activity at the neighboring frequencies (reference, **R**).

For each participant, 82 spatial filters were constructed (separate for each flicker frequency and hemifield, but collapsed across “attend flicker” and “ignore flicker” conditions), because: (1) SSVEP topographies depend on the location of the flickering stimuli in the visual field, and for lateralized stimuli are maximal over the contralateral hemisphere (Kim et al., 2007; Morgan et al., 1996; Spaak et al., 2014; Figure 3); (2) different frequency SSVEPs have different sources and therefore different scalp projections (Heinrichs-Graham and Wilson, 2012; Lithari et al., 2016), and (3) SSVEP topographies may differ across participants due to anatomical differences.

Each spatial filter was designed as described further. First, channels with high variance EEG signal containing EMG noise (mostly frontal and temporal) were identified using Fieldtrip *ft_rejectvisual* function (Oostenveld et al., 2011; http://www.ru.nl/neuroimaging/fieldtrip/), and were excluded due to their negative impact on the accuracy of RESS spatiotemporal filters (Cohen and Gulbinaite, 2017). Second, condition-specific single-trial data were concatenated and temporally filtered using narrow-band Gaussian filters: (1) filter centered at the flicker frequency *f* with full-width at half-maximum (FWHM) = 0.6 Hz; (2) filter centered at *f*-1 Hz with FWHM = 2 Hz; (3) filter centered at *f*+1 Hz with FWHM = 2 Hz. Data filtered at the flicker frequency and the neighboring frequencies are further referred to as “signal” (**S**) and “reference” (**R**), respectively. Narrow-band filtering was performed in the frequency domain by multiplying the Fourier coefficients of the channel time series with frequency-domain Gaussian filter:

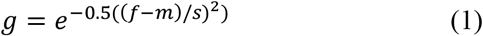

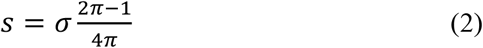

where *f* is a vector of frequencies from the Fourier transform of the time series signal, *m* is the peak frequency of the filter, and *σ* is the desired spectral FWHM. After frequency-wise multiplication, the inverse Fourier transform was applied to obtain the narrowband filtered time series signal.

Third, temporally-filtered data from 1500 to 7000 ms (relative to the cue onset) were used to compute channel covariance matrices (two R matrices that were averaged and one S matrix). Flicker-onset and offset elicits strong visual responses, which affect the quality of the spatial filter and thus were excluded from the analyses (Cohen and Gulbinaite, 2017). To increase the robustness of the spatial filters, we applied a 1% shrinkage regularization to the **R** covariance matrix. Shrinkage regularization involves adding a percent of the average eigenvalues onto the diagonals of the **R** covariance matrix. This reduces the influence of noise on the resulting eigendecomposition. Fourth, generalized eigenvalue decomposition (MATLAB function *eig*) on the covariance matrices **S** and **R** was used to construct spatial filters (**W**):

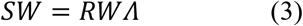

where **W** is the matrix of eigenvectors and **Λ** is a diagonal matrix of eigenvalues. The eigenvectors (column vectors with values representing electrode weights) were used to obtain RESS component time series (eigenvector multiplied by the original unfiltered single-trial time series). Although the first RESS component associated with the highest eigenvalue typically has the highest SNR at the frequency of interest (i.e., EEG responses at the frequency that the spatial filter is designed to maximize), power spectra of RESS components were expressed in SNR units, and the RESS component with the highest SNR at the flicker frequency was automatically selected for subsequent analyses. Of 1552 RESS components (1552 = 19 participants x 41 flicker frequency x 2 attention conditions), the first RESS component was selected for subsequent analyses in 94.52% of cases, the second RESS component in 4.12% of cases, and third RESS component in 1.35% of cases. Thus, for each trial we analyzed separate hemifield- and frequency-specific RESS component time series. Topographical maps of spatial filters (Y) were obtained by left-multiplying the eigenvectors by the **S** covariance matrix:

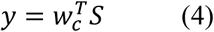

where *w* is the *c*^th^ column in **W** matrix. Maps were normalized prior to averaging across participants (Figure 3A and C, Figure 5D, and Figure 7B).

Although flicker frequency-specific spatial filters increase topographical specificity of SSVEP signal, to determine how important this topographical specificity is we constructed common spatial filters using SSVEP responses to all flicker frequencies. Thus for each participant we had 2 common spatial filters (separate for each hemifield but collapsed across “attend flicker” and “ignore flicker” conditions). For construction of common spatial filters, we used RESS as described above. The covariance matrix S was defined as average of covariance matrices temporally filtered at the stimulus frequency (41 matrices), and the covariance matrix **R** was defined as average of covariance matrices temporally filtered at the neighboring frequencies (82 matrices). All temporal filter parameters were kept the same as in frequency-specific SSVEP responses.

For construction of RESS spatial filters specific to harmonic responses (2*f*, 3*f*, 4*f*, 5*f*, and 6*f*) we followed the same procedure as for the SSVEPs at the stimulation frequency (described above). We constructed separate spatial filters for each harmonic of the stimulus frequency because harmonic SSVEP responses have different cortical foci, and thus different topographical distributions (Heinrichs-Graham and Wilson, 2012; Pastor et al., 2007).

### SSVEP analysis: Total SSVEP response

Although typically SSVEP power is determined by applying a Fourier transform to trial-average waveforms (Kim et al., 2007; Muller et al., 1998b), such approach allows to extract only the phase-locked part of the SSVEP signal. The non-phase locked part of the SSVEP signal contains attention-related modulation of SSVEP amplitude over time and is represented by the side-bands around the flicker frequency in the power spectrum (Nunez and Srinivasan, 2006). To preserve the non-phase locked part of the signal, we applied a Fourier transform on single-trial data in a 1500–7000 ms time window (relative to the flicker onset). To obtain equally good resolution for all flicker frequencies (0.1 Hz), the exact time window for FFT was adjusted by zero-padding the data. The absolute value of FFT coefficients was squared and averaged across trials. To facilitate comparison across different flicker frequencies and to diminish the effect of endogenous oscillations on estimates of SSVEP response, SSVEP power values were converted into SNR units (Norcia et al., 2015). SNR was computed as the power at the frequency of interest divided by the average power at the neighboring frequencies (+-1 Hz, excluding 0.5 Hz around the frequency of interest, which respectively corresponds to 10 and 5 frequency bins at 0.1 Hz frequency resolution).

For the time-course analysis of SSVEP amplitudes (Figure 4B), the frequency-specific RESS time series was filtered using a narrow band-pass Gaussian filter, with FWHM = 1 Hz to ensure high frequency specificity. After filtering the data, the instantaneous SSVEP amplitude was extracted using a Hilbert transform. Trial average amplitude was baseline normalized to the pre-cue period of −600 to −200 ms by computing percent change of the amplitude at each time point relative to the average amplitude of the baseline period. This normalization procedure allows comparison across flicker frequencies and eliminates scale differences between individuals. Thereafter, amplitude time series were aligned separately to individual alpha and gamma resonance frequencies, such that for each subject resonance frequency was at zero.

### SSVEP analysis: Phase-locked part of SSVEP response

To investigate the contribution of lateralized endogenous alpha to the reversed attentional modulation pattern for the alpha-band SSVEPs, we analyzed only the phase-locked part of the SSVEP signal. The small number of trials per experimental condition prevented us from using a trial-average approach to extract only the phase-locked part of the SSVEP signal (Kim et al., 2007; Muller et al., 1998b). Therefore, we used a sliding window approach by time-locking the RESS (and electrode) time series to stimulus luminance peaks (Muller et al., 1998a; Wieser et al., 2016). For each flicker frequency, the sliding window spanned 4 cycles, and was shifted in steps of one SSVEP cycle. We then applied FFT to the window-averaged waveform and extracted SSVEP power by squaring the absolute value of FFT coefficients. Based on previous reports of increased SSVEP phase coherence for attended vs. ignored stimuli (Kashiwase et al., 2012; Keitel et al., 2017; Kim et al., 2007), we also computed phase consistency of SSVEPs by averaging *e^ik^*, where *k* is the phase angle of the complex FFT coefficients. The absolute value of the result is also known as phase-locking factor or inter-trial phase clustering (Lachaux et al., 1999; Makeig et al., 2004).

### Alpha-band power (8-13 Hz) analyses

To evaluate α-band power changes related to the shifts and maintenance of spatial attention separately from attentional modulation of SSVEPs, only 15-80 Hz flicker conditions were included in the α-power analyses, meaning that SSVEPs (1*f* and harmonically-related responses) in the alpha-band were excluded. We performed two sets of analyses. First, to make alpha-band power results comparable to SSVEP analyses that were done in the frequency domain, we estimated alpha power using short-time FFT in the same time window (1500-7000 ms), with a 500-ms sliding window, and steps of 25 ms (i.e. 95% overlap between successive time segments). Second, we estimated alpha power changes over time by convolving stimulus-locked single-trial data from all electrodes with complex Morlet wavelets, defined as:

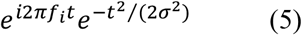

where *t* is time, *f_i_* is frequency which ranged from 2 to 60 Hz in 50 logarithmically spaced steps, and *σ* is the width of each frequency band defined as *n*/(*2πfi*), where *n* is a number of wavelet cycles that varied from 3 to 15 in logarithmically spaced steps. Thereafter, we computed instantaneous power by taking the square of the complex convolution result, averaging over trials, and normalizing power values by converting to decibel scale relative to the pre-stimulus time window (−500 – −200 ms, where 0 is the onset of the cue).

### Fluctuations in instantaneous SSVEP response frequency

We investigated the stability of SSVEP phase by estimating fluctuations in instantaneous frequency. Instantaneous frequency is defined as the first temporal derivative of the phase time series (Cohen, 2014b). Condition-specific single-trial phase time series was obtained by temporally filtering RESS component time series using a Gaussian filter centered at the flicker frequency (FWHM = 4 Hz) and applying the Hilbert transform. The instantaneous frequency was computed by taking the first derivative of the unwrapped phase time series and converted to Hz by multiplying the result by the sampling rate and dividing by 2*π* (Cohen, 2014b):

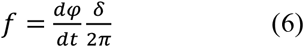

where *φ* is the time series of unwrapped instantaneous phase angles, and *δ* is the sampling rate.

Spikes in the EEG time series appear as sudden changes in the phase time series (“phase slips”), which appear as sudden increases or decreases in instantaneous frequency but are not physiologically plausible. These abrupt frequency changes were attenuated using a two-step procedure. First, for each trial extreme outliers (exceeding 100 Hz and falling outside 99 percentile of frequency values in that trial) were identified and replaced by the average of neighboring instantaneous frequency values. Second, remaining outliers were replaced by the median of instantaneous frequency estimates in a +-20 ms window surrounding the outlier. We then concatenated all the trials, separately for each flicker frequency and attention condition, and computed a standard deviation of instantaneous SSVEP frequency changes over time. To estimate fluctuations of instantaneous frequency that can be expected in the absence of flicker (null hypothesis), we repeated the same procedure using trials in which neither flicker frequency nor harmonically related responses were present, as well as trials that had flicker frequency within the boundaries of the frequency response of the Gaussian filter kernel of interest.

#### Statistical analyses

##### Behavioral data analyses

We performed a repeated-measures ANOVA with trial validity (valid and invalid), stimulus background (flicker and static), and hemifield (left and right) as within-subject factors. Given that the frequency window for the QUEST procedure was set to +-5 frequencies surrounding the frequency of interest, trial-average luminance values for each flicker frequency condition were smoothed using a running average of ±5 neighboring flicker frequency conditions prior to trial-average statistical analyses of blue LED luminance values (target stimuli) as a function of flicker frequency and target stimulus background (Figure 2B, C). Note that luminance values of blue LEDs were also dynamically adjusted during the experiment to keep the performance on valid trials at 80% accuracy using QUEST procedure (taking into account performance over the previous 40 trials in that flicker frequency ±5 frequencies surrounding the frequency of interest, done separately for static and flickering background). Statistical comparison of target luminance values for static and flickering background (collapsed across hemifields) was performed using a nonparametric permutation testing procedure correcting for multiple comparisons across frequencies using cluster-level correction (Maris and Oostenveld, 2007).

**Figure 2.**
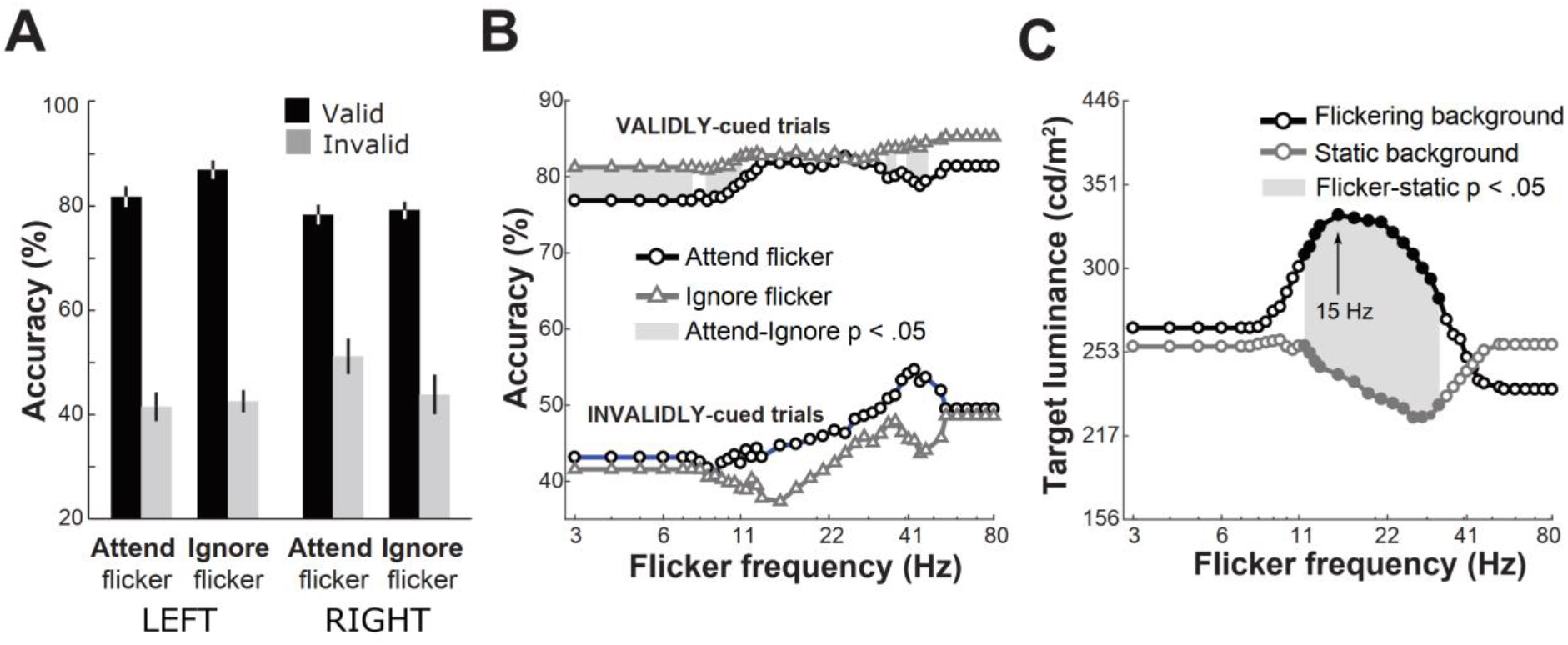
Behavioral results. (A) Performance accuracy as a function of trial validity (valid trials = report the number of blue flashes on the cued side, invalid trials = report the number of blue flashes on the uncued side), the background against which the to-be-detected blue stimulus was presented (attend flicker = flickering background, ignore flicker = static background), and hemifield (right, left). Error bars denote SEM. (B) Performance accuracy as a function of flicker frequency and state of the background against which blue stimuli were presented. Top two lines fluctuating around 80% represent performance on validly-cued trials, the bottom two lines – invalidly-cued trials. Gray shaded areas represent frequencies at which target detection accuracy for static vs. flickering background statistically significantly differed (p < .05, uncorrected for multiple comparisons across frequencies). (C) Average target stimulus luminance as a function of flicker frequency. Note that luminance values of blue LEDs were dynamically adjusted for static and flickering background separately using a QUEST procedure considering performance over the previous 40 trials and flicker frequency ±5 frequencies surrounding the frequency of interest. Therefore, for the lowest and highest 6 flicker frequencies (3-7.5 Hz and 55-80 Hz) the average luminance values are the same. Gray shaded area represents frequencies at which target luminance for static vs. flickering background statistically significantly differed (p < .05, uncorrected for multiple comparisons across frequencies). Filled-in data points represent the same contrast corrected for multiple comparisons across frequencies.

**Figure 3.**
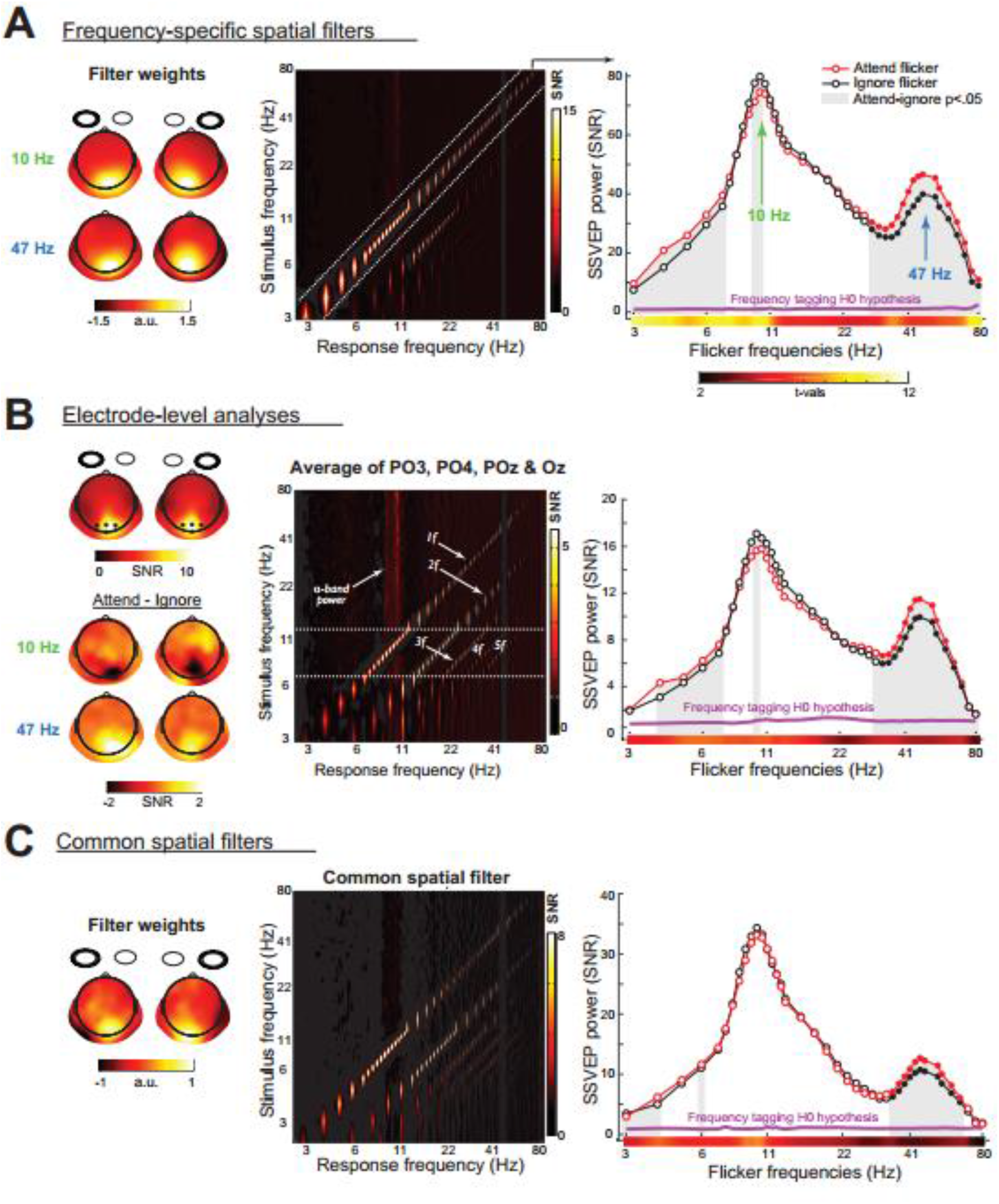
Summary of EEG results. (A) SSVEP responses isolated using frequency-specific spatial filters. Left panel: Example topographical maps of spatial filters for 10 Hz and 47 Hz flicker conditions. Middle panel: Subject-average response amplitude expressed in SNR units (x-axis) plotted as a function of stimulus flicker frequency (y-axis). The brightest colors along the diagonal indicate that maximal SSVEP responses matched the stimulus frequency. Note the relative suppression of harmonic responses and endogenous alpha compared to electrode-level spectral response profile in B. Right-panel: SSVEP power at all 41 flicker frequencies, plotted separately for “attend flicker” (red line) and “ignore flicker” (black line) conditions (collapsed across left and right hemifield flicker conditions). Note the resonance peaks in response to alpha- (~10 Hz) and gamma-band flicker (~ 47 Hz). Magenta line represents SSVEP amplitude in trials when neither flicker nor harmonic responses were present at that frequency (the null hypothesis). Statistically significant SSVEP amplitudes were observed for all tested frequencies (p < .001, corrected for multiple comparisons across frequencies using cluster-based permutation testing). The actual t-values for this comparison are color-coded on the x-axis. Grey shaded areas represent statistically significant SSVEP amplitude differences between “attend flicker” and “ignore flicker” conditions (uncorrected for multiple comparison across flicker frequencies). Filled-in data points represent the same contrast corrected for multiple comparisons across frequencies. (B) Electrode-level SSVEP analyses using occipito-parietal electrodes POz, Pz, PO3, PO4, which were selected based on the condition-average topographical maps (left panel). Below, contrast between “attend flicker” and “ignore flicker” conditions at resonance frequencies (10 Hz and 47 Hz) depicting negative attentional modulation for alpha-band flicker, and positive – for gamma-band flicker. Topographically, attentional modulation was maximal contralaterally to the flickering stimulus. (C) SSVEP responses isolated using common spatial filters, which disregard the topographical specificity of SSVEP response at different flicker frequencies. Note the differences in SNR values between A, B, and C.

##### Statistical analyses of SSVEPs

Given that spatial filters are specifically designed to maximize power at the flicker frequency while suppressing activity at the neighboring frequencies, SNR > 1 values can be expected even in the random data. To determine SNR values (“noise SNR”) that can be expected due to overfitting and obtain a statistical estimate of robustness of flicker-induced EEG activity, for each flicker frequency and hemifield we created RESS spatial filters using trials, in which neither flicker frequency nor harmonically related responses were present (e.g. for estimation of noise SNR for 20 Hz flicker, all but 5 Hz, 10 Hz, and 20 Hz flicker trials were included). “Noise” RESS spatial filters were used to obtain “noise” RESS component time series, which were further used to compute noise SNR expected under the null hypothesis following the same procedure as for the SSVEP amplitude.

To compare the results obtained using the RESS procedure to electrode-level analyses – an approach more commonly used in the SSVEP literature (Kashiwase et al., 2012; Keitel et al., 2013; Morgan et al., 1996; Muller et al., 1998a; Toffanin et al., 2009) – we selected a subset of electrodes that showed maximum SSVEP power (collapsed across “attend flicker” and “ignore flicker” conditions, and across flicker frequencies). For left-hemifield flicker trials PO4, POz and Oz electrodes were selected, whereas PO3, POz, Oz showed maxima for the right-hemifield flicker trials (Figure 3B, left panel). For statistical analyses of SSVEP amplitude, we obtained noise SNR values by averaging power at the flicker frequency of interest from the trials, in which neither flicker frequency nor harmonically related responses were present.

Prior to trial-average statistical analyses, SNR values for each flicker frequency were smoothed using a running average of ±2 neighboring flicker frequency conditions due to the small number of trials per flicker frequency (N=12). Smoothing procedure was done at the subject level, separately for each condition (attend flicker and ignore flicker) and hemifield (left and right). Statistical significance of the effects of flicker frequency on the SSVEP amplitude was evaluated by comparing the observed SNR value to that of noise SNR expected under the null hypothesis using a nonparametric permutation testing procedure correcting for multiple comparisons across frequencies using cluster-level correction (Maris and Oostenveld, 2007). The same statistical procedure was performed when evaluating attentional modulation across different flicker frequencies, i.e. comparing SNR spectra for attend flicker and ignore flicker conditions.

##### Statistical analyses of alpha power

Statistical significance of alpha-band lateralization in attend vs. ignore flicker conditions (Figure 5D) was evaluated using the following nonparametric permutation testing procedure. First, real t-values were obtained at each pixel of the topographical map (electrode-level alpha-band power extrapolated using Matlab plotting function *contourf*) by performing a one-sample t-test comparing the condition difference against 0. Second, a null hypothesis distribution of t-values (permuted t-values) was generated by swapping condition labels and rotating the topographies by 90° for a random number of subjects, and re-computing t-values. This procedure was repeated 1000 times. Rotation of topographies prevents from obtaining artificially high t-values at channels that have very small variance across subjects. Third, the real t-values were z-scored using the mean and SD of permuted t-values. Fourth, power differences between conditions were considered statistically significant if the z-scored t-value at each pixel in the topographical map was larger than 4.2649 (*p* = .0001). Finally, cluster-based correction was applied to correct for multiple comparisons over pixels. The null hypothesis distribution of cluster sizes was obtained by thresholding topographical maps from each of the 1000 iterations at *p* = .0001, and storing the maximum cluster size value (sum of t-values within a cluster). Clusters of contiguous pixels in the real t-value topographical maps were considered significant if the size of the cluster was bigger than expected under the null hypothesis. To obtain more stable estimates from permutation testing, we ran a “meta-permutation test” by repeating pixel-level and cluster-level permutation procedure 20 times. Thus, the average of 20 real z-scored t-values and the average of 20 cluster thresholds were used.

## RESULTS

### Behavioral results

Based on numerous reports on behavioral advantages of valid spatial cues in spatial attention tasks (for a review, see Carrasco 2011), we expected participants to be more accurate in reporting the number of blue flashes on the cued side (valid trials) as compared to the uncued side (invalid trials). For behavioral data analysis, we performed a repeated-measures ANOVA with trial validity (valid and invalid), stimulus background (flicker and static), and hemifield (left and right) as within-subject factors. Behavioral results are summarized in Figure 2.

Despite the presence of flicker, we observed a typical effect of cue validity: Participants were more accurate on valid vs. invalid trials (*F*_(1,18)_ = 122.55, *p* < .001, η = .872). Blue flashes presented in the flickering vs. static background were equally well detected (no main effect of flicker, *F*_(1,18)_ = 0.005, *p* = .946, η < .001), and we found no differences in detection performance between right- and left-hemifield stimuli (no main effect of hemifield, *F*_(1,18)_ = 0.025, *p* = .875, η = .001). However, there was a significant interaction between hemifield and stimulus background state (*F*_(1,18)_ = 9.44, *p* = .007, η = .344), with more accurate performance for left-hemifield targets presented on static vs. flickering background and the opposite pattern for the right-hemifield targets. The validity effect also differed between hemifields (*F*_(1,18)_ = 12.59, *p* = .002, η = .412). Follow-up analyses revealed that accuracy was higher for left-hemifield targets on valid trials (*t*(18) = 4.68, *p* < .001), and there was a tendency for higher accuracy for right-hemifield stimuli on invalid trials (*t*(18) = −1.98, *p* = .063), meaning that more distractors on the left-hemifield were noticed. Such accuracy differences between hemifields could reflect a well-documented attentional bias towards the left hemifield (Buschman and Kastner, 2015).

We also analyzed performance accuracy as a function of flicker frequency and state of the background against which blue stimuli were presented (static vs. flickering white LEDs). Given that target luminance values were dynamically adjusted to keep performance at 80% accuracy level taking into account flicker frequency, and using separate QUEST staircases for attend flicker and ignore flicker conditions but the same staircase for valid and invalid trials, deviations from the expected 80% accuracy are possible and of interest (see Methods section for further details). As depicted in Figure 2B, performance accuracy varied as a function of flicker frequency. For validly cued trials, statistically significant differences between target detection on flickering vs. static background were present during theta, alpha and gamma flicker (p < .05 3-7.5 Hz, 8.5-11, 35-37 Hz, and 41-47 Hz, uncorrected for multiple comparisons across frequencies). Although overall distractors presented on the static background were more often noticed (blue line for invalidly-cued trials), the effect was not statistically different for any flicker frequency.

Target stimulus visibility interacted with flicker in a frequency-dependent manner, with significant target luminance differences between flickering and static background for flicker frequencies centered on the beta band (11.5-33 Hz corrected for multiple comparisons across frequencies). On average, 15 Hz background flicker required the highest luminance of the blue targets to be detected with 80% accuracy. Importantly, flicker frequency affecting target detection (Figure 2B) and brightness perception (Figure 2C) did not overlap. Taken together, theta, alpha and gamma flicker had cognitive effects, whereas beta flicker affected brightness perception.

#### EEG results

##### Resonance response to alpha- and gamma-band flicker

Following previously reported SSVEP amplitude variability as a function of stimulation frequency, with resonance peaks consistently reported around ~10 Hz (Fedotchev et al., 1990; Herrmann, 2001; Lazarev et al., 2001; Regan, 1977), and in some cases around ~15 Hz (13-25 Hz range, Herrmann, 2001; Pastor et al., 2007) and 40-55 Hz (Herrmann, 2001; Regan, 1968), we expected to observe local maxima in SSVEP responses. Contrary to previously reported qualitative descriptions of resonance peaks, we took a rigorous quantitative approach. We compared the general pattern of SSVEP responses to different flicker frequencies using three complementary analyses: (1) a traditional SSVEP data analysis approach using occipito-parietal electrodes (POz, Oz, PO3, PO4) that showed maximal response across all flicker frequencies (Figure 3B); (2) frequency-specific spatial filters making use of topographical differences of SSVEPs generated in response to each flicker frequency (Figure 3A); (3) common spatial filters, which were constructed ignoring topography of frequency-specific SSVEPs (Figure 3C).

As expected based on retinotopic organization of the primary visual cortex, lateralized flickering stimuli elicited maximal responses at occipito-parietal electrodes contralateral to the flickering stimulus location (Figure 3, left panels). There were also topographical differences in maximal responses for low vs. high flicker frequencies: More posterior and lateralized peak response to 10 Hz as compared to more anterior and central peak in response to 47 Hz flicker (see topographical representation of RESS spatial filter weights in Figure 3A, left panel). Given these subtle but important topographical differences, the use of subject- and frequency-specific spatial filters allowed to maximize the SNR of the narrow-band SSVEP response (compare SNR values in Figure 3 across middle panels).

The spectral response profile showed the most prominent peaks along the diagonal using all three analyses approaches (electrode-level, frequency-specific and common spatial filters), reflecting high frequency specificity of SSVEPs (Figure 3, middle panels). Response peaks were also present off the diagonal at multiple harmonics of the stimulus frequency (2*f*, 3*f*, 4*f*, etc.). As expected, harmonic responses were more pronounced for electrode-level and common-spatial filter analyses because frequency-specific RESS spatial filters suppress the signal outside the narrow-band flicker frequency of interest. However, given that the RESS method is based not only on temporal (flicker frequency) but also spatial (topographical distribution) characteristics of SSVEPs, the presence of harmonic responses off the diagonal in Figure 3A middle panel are consistent with the reports of partially spatially-overlapping cortical foci of the fundamental and higher harmonic responses (Heinrichs-Graham and Wilson, 2012; Pastor et al., 2007).

Statistically significant SSVEP responses were present up to 80 Hz flicker, with the strongest response, or *resonance*, in response to alpha (~10 Hz) and gamma (~47 Hz) flicker. In Figure 4, subject-specific spectral response profiles normalized to the SSVEP amplitude at each participant’s alpha and gamma resonance frequencies reveal variability of resonance peaks across participants, and illustrate that higher resonance frequency in alpha is not necessarily associated with higher resonance frequency in gamma (*r*_Pearson(17)_ = - .121, *p*_two-tailed_ = .621). Time course analysis of SSVEP amplitude revealed that SSVEPs around alpha and gamma resonance frequencies ramped up earlier after the flicker onset and persisted longer after the flicker offset (Figure 4B). These two features – early onset and reverberations after offset – can be taken as partial evidence for the entrainment of endogenous oscillations by external rhythmic input (Thut et al., 2011). Consistent with previous reports (Herrmann, 2001; Pastor et al., 2007), a resonance peak in the beta band was also present for some participants. However, due to variability in peak frequency across participants (15-23 Hz) and relatively small amplitude, the grand-average spectral response profile in Figure 3 right column does not contain a prominent peak in responses to beta-band flicker.

**Figure 4.**
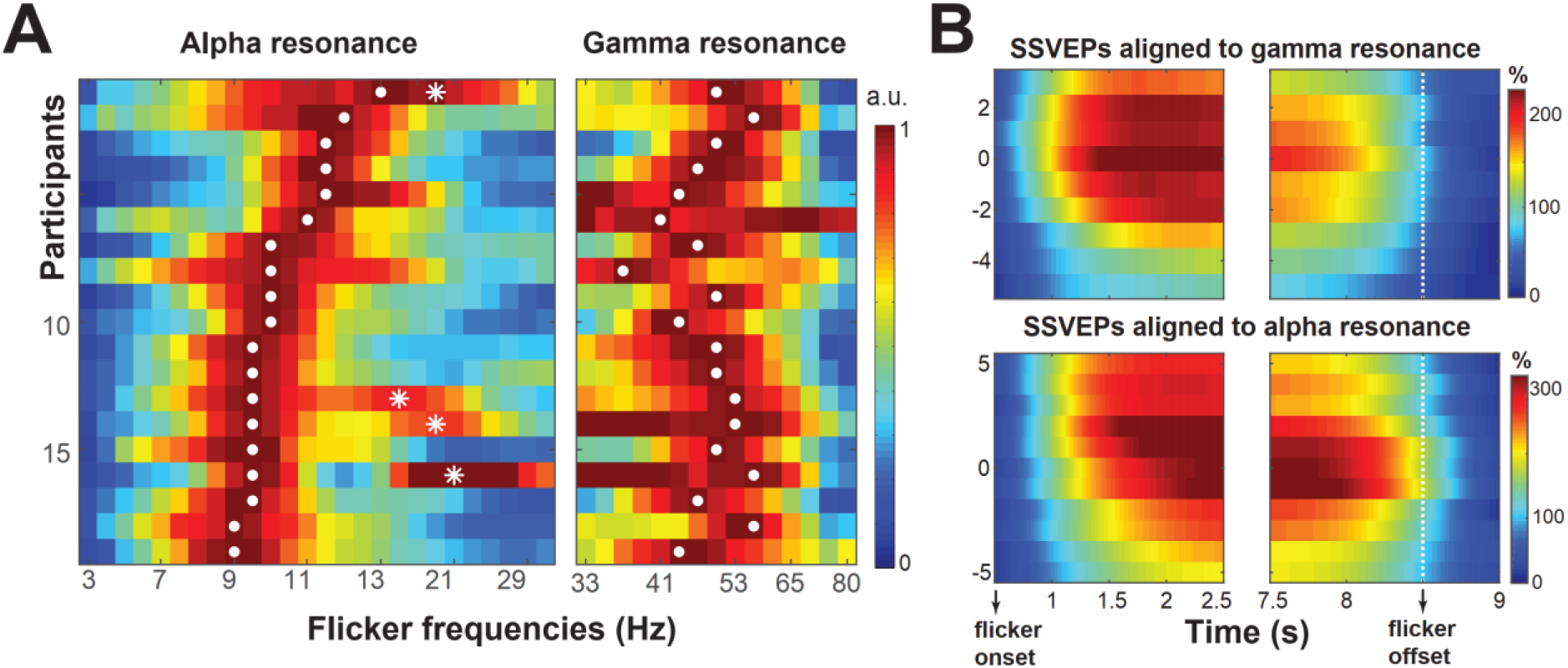
Subject-specific resonance frequencies for alpha- and gamma-band flicker. (A) Subject-specific spectral response profile normalized to the SSVEP amplitude at each participant’s resonance response frequency in alpha (left panel), and gamma (right panel). Separate normalization performed due to amplitude differences between alpha and gamma resonance peaks. Each row represents the spectral response profile of one participant, with the same order across the two panels. Color represents normalized SNR values. White dots represent the individual’s resonance frequencies in alpha and gamma, stars represent resonance frequencies in the beta band that were present for some participants. (B) SSVEP time courses averaged across participants, with 0 Hz representing each participants gamma (top panels) and alpha (bottom panels) resonance frequency. The time courses are expressed as % change relative to pre-cue period (−600 – 200 ms), where cue onset is 0 s. Note the early ramping up of SSVEP amplitude around the resonance frequencies following the flicker onset, and longer persistence of increased amplitude around the flicker offset.

We estimated statistical robustness of flicker-related activity by comparing the observed SSVEP SNR values to those of noise SNR values obtained using trials in which neither flicker frequency nor harmonically related SSVEP responses were present (see Methods section for details). For all tested frequencies and all three analysis approaches (Figure 3A, B, C), the observed SNR was significantly higher than noise SNR (p < .001, corrected for multiple comparisons across frequencies; see t-value statistics of this comparison as a color-code on the x-axis in Figure 3, right column).

### Attention increases gamma-band and decreases alpha-band SSVEP amplitude

All three analysis approaches consistently revealed that attentional modulation varied in size and direction of the effect (see Figure 3, right column). In response to theta-band flicker (3-7 Hz, corrected for multiple comparison across flicker frequencies), SSVEP amplitude was significantly higher for “attend flicker” vs. “ignore flicker” (positive effect). Attention also positively modulated gamma-band SSVEP amplitude (29-80 Hz frequency range, corrected for multiple comparisons across flicker frequencies), with on average 20.22% (SD = 11.14%) increase in SSVEP amplitude at subject-specific gamma resonance frequency. Alpha-band SSVEPs (8-13 Hz), however, showed negative attentional modulation across the entire alpha band and significantly negative attentional modulation for 9.5 and 10 Hz flicker (uncorrected for multiple comparison across flicker frequencies), with on average 6.62% (SD = 11.10%) decrease in SSVEP amplitude at the subject-specific alpha resonance frequency (see Figure 5D for a direct contrast between “attend flicker” and “ignore flicker” conditions across participants and effect size).

**Figure 5.**
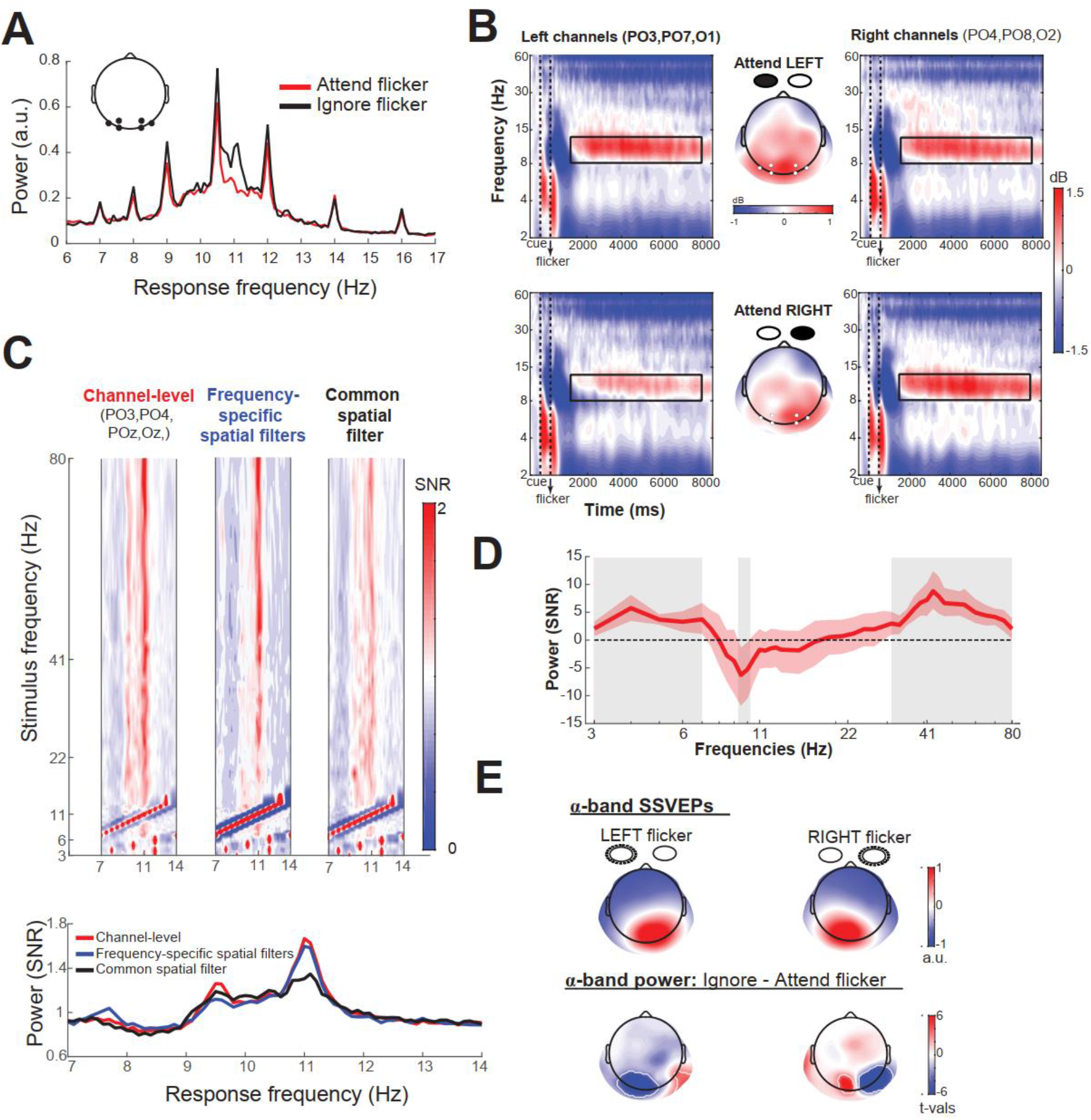
Spatial overlap between endogenous alpha-band oscillations and alpha-band SSVEPs. (A) Grand-average power spectra from several alpha-band flicker conditions (selected for illustrative purposes). Each line represents the grand-average across occipitoparietal channels contralateral to the flicker side (PO3, PO7, O1 for the right-flicker trials; PO4, PO8, O2 for the left-flicker trials) illustrating co-occurrence of SSVEPs (peaks at 7, 8, 9, 10.5, and 12 Hz) and endogenous alpha-band oscillations (general increase of power around ~10 Hz). For illustrative purposes, only a subset of flicker frequencies is plotted. (B) Time-frequency representations of spatial attention dynamics over posterior channels, separated for two attention conditions: attend left (top panels) and attend right (bottom panels). Note the changes in alpha-band oscillatory response during the trial: Decrease in alpha-band power in response to cue and flicker onset, followed by the sustained increase in alpha power (relative to the pre-cue baseline period). Topographical maps show mean alpha-band power distribution over the scalp in time-frequency windows outlined by black rectangles: 8-13 Hz, 1500-8000 ms post-cue. (C) Zoomed-in version of spectral profiles depicted in Figure 3 middle panels, highlighting the differences in the amount of endogenous alpha-band power captured by three different SSVEP data analyses approaches: channel-level, flicker-frequency specific spatial filters, flicker frequency non-specific (common) spatial filters. Below: 2D version of the plot in panel C, collapsed across 15-80 Hz flicker conditions to allow a more direct comparison. (D) SSVEP amplitude difference between “attend flicker” and “ignore flicker” conditions (same data as plotted in Figure 3A right-most panel). Red shaded areas represent 95% percentile confidence intervals around the mean, estimated using a bootstrapping procedure (N = 10000 iterations, resampling with replacement). Grey shaded areas represent statistically significant SSVEP amplitude differences between “attend flicker” and “ignore flicker” conditions (uncorrected for multiple comparison across flicker frequencies). (E) Top: Subject-average topographical maps of frequency-specific RESS spatial filters (8-13 Hz flicker frequencies). Bottom: Alpha-band power (8-13 Hz) difference between “ignore flicker” and “attend flicker” conditions (plotted only for 15-80 Hz flicker conditions). White areas outline the regions in which the two attentional conditions significantly differed at p < .0001 two-tailed (corrected for multiple comparisons across pixels using cluster-based permutation testing).

As can be seen in Figure 5A, subject-average raw power spectra at occipito-parietal channels contain both responses to alpha-band flicker (narrow peaks at 7, 8, 9, 10.5, and 12 Hz flicker frequencies, selected for illustrative purposes), as well as a general increase in power around ~10 Hz (endogenous alpha-band oscillations). Thus, reversed attentional modulation could reflect a linear superposition of alpha-band oscillations implicated in visuospatial attention and alpha-band SSVEPs at the scalp level, or an interaction between endogenous alpha oscillations and alpha-band flicker. Although the SSVEP time-course analysis at resonance frequencies provided partial support for the latter alternative (Figure 4B), we next analyzed the temporal dynamics of endogenous alpha and the spatial distribution of attention effects on SSVEPs and endogenous alpha.

### Attention-related changes of endogenous alpha-band oscillations

Temporal dynamics of alpha power (collapsed between left-flicker and right-flicker trials) during the trial relative to the pre-cue baseline period followed the previously reported two stages (Rihs et al., 2009; Worden et al., 2000): Early decrease in alpha power related to orienting of spatial attention (endogenous cue-related and exogenous flicker-onset-related), and late increase in alpha power, reflecting sustained attention (Figure 5B). We focused on the late part, as it coincided with the time window of SSVEPs. As expected based on numerous previous findings on alpha lateralization in covert spatial attention tasks (for a review, see Frey et al., 2015), alpha power was enhanced over the occipital areas representing the to-be-ignored part of the visual space (i.e. contralaterally to the distractors) relative to the areas representing the to-be-attended space. Therefore, not surprisingly, the electrode-level spectral response profile contained a prominent alpha-band peak (~10 Hz), which was present regardless of the flicker frequency (Figure 5C left).

To compare spatial distribution of attention effects on alpha-band power and alpha-band SSVEPs directly, we performed frequency-domain analyses of alpha power using short-time FFT in the 1500-7000 ms using only conditions that did not contain flicker or harmonically-related responses in the alpha band (i.e. 15-80 Hz flicker conditions). Although exclusion of SSVEP frequencies in the FFT spectra has been used to separate SSVEP from endogenous alpha oscillations (Ding et al., 2006; Kelly et al., 2006), such approach assumes SSVEP to be a perfectly stationary narrow-band signal. This assumption, however, does not take into account that the amplitude modulation of SSVEP signal related to fluctuations in attention (which is a relatively slow process) is captured by the side-bands around the stimulation frequency (Cohen and Gulbinaite, 2017; Nunez and Srinivasan, 2006). Thus, exclusion of SSVEP frequencies from the FFT spectrum does not allow a clean separation of SSVEPs from endogenous activity.

For both left- and right-flicker trials, we observed statistically significant alpha power lateralization (p < .0001 two-tailed, corrected for multiple comparisons across pixels using cluster-based permutation testing; Figure 5B): Stronger alpha power over occipital areas representing the to-be-ignored part of the visual space (i.e. contralaterally to the to-be-ignored flicker). Note that to facilitate interpretation of reversed attention effects on SSVEP amplitude (Figure 5D), we contrasted “ignore flicker” and “attend flicker” conditions. As can be seen in Figure 5E, the maximal response to alpha-band flicker and attention effects on endogenous alpha are spatially overlapping, suggesting that superposition of the two signals could explain the observed negative attentional modulation of alpha-band SSVEPs.

### Positive effect of attention on phase-locked part of alpha-band SSVEPs

Endogenous alpha-band power was less pronounced in the spectral response profile derived using frequency-specific spatial filters, and even less so using common spatial filters (Figure 5C, line plots). This result indicates that RESS spatiotemporal filters captured part of the endogenous alpha signal. Partial rather than complete separation of SSVEPs from endogenous alpha oscillations reflects spatial overlap of the two signals, and possible temporal interaction. Therefore, we next exploited differences in temporal dynamics between the two signals as a means to estimate the contribution of endogenous alpha to the reversed effects of attention on alpha-band SSVEPs. We reasoned that attention-related changes in endogenous alpha rhythm would have a random phase with respect to the flickering stimulus, and thus by time-locking EEG time series to stimulus luminance peaks, we would eliminate or at least reduce the contribution of endogenous signals that are non-phase-locked to the driving stimulus. We used a sliding window approach (4-cycle window at each flicker frequency shifted in steps of one cycle), and analyzed power differences between “attend flicker” and “ignore flicker” conditions, as well as phase alignment between stimulus and response waveforms by computing phase clustering across sliding windows (a procedure mathematically identical to computation of inter-trial phase clustering, ITPC; Cohen, 2014a).

The amplitude of the phase-locked part of the SSVEP signal varied as a function of frequency, with a local peak in alpha (Figure 6A). Note that here SSVEP response is reported without conversion to SNR units due to low frequency resolution resulting from short sliding window size. Attentional modulation of the phase-locked part of the SSVEP signal was significantly positive for the theta- and also numerically positive for the alpha-band SSVEPs (Figure 6A). Analyses using 6 and 8-cycle sliding windows showed nearly identical patterns of results.

**Figure 6.**
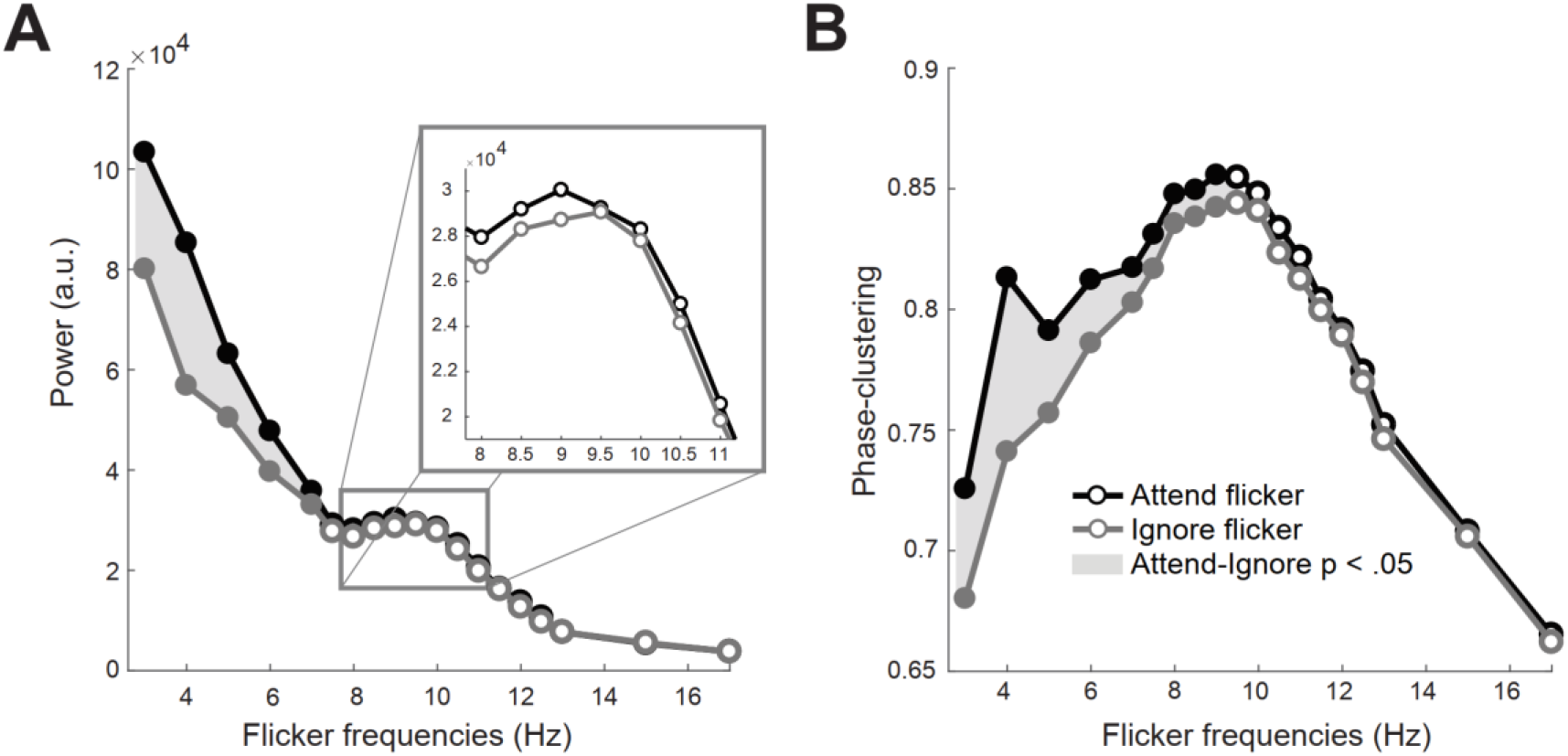
Attentional modulation of phase-locked SSVEPs. (A) Raw SSVEP power as a function of flicker frequency plotted separately for “attend flicker” (black line) and “ignore flicker” (grey line) conditions (collapsed across left and right hemifield flicker conditions). Conversion to SNR units could not be performed due to low frequency resolution imposed by the sliding window approach. (B) Phase-clustering as a function of flicker frequency. Conventions are the same as in Figure 3.

Phase-clustering analysis revealed a stronger phase-alignment of SSVEP waveform to alpha-band driving stimuli (peak ~ 10 Hz), indicating that the resonance in SSVEP amplitude in response to alpha-band flicker results from more stable SSVEP phase (Figure 6B). Moreover, phase clustering was statistically significantly stronger for “attend flicker” than for “ignore flicker” conditions for 3-9 Hz frequencies (corrected for multiple comparisons across frequencies). This finding corroborates previous reports on increased inter-trial phase clustering of SSVEPs for attended vs. ignored flicker stimuli (Kashiwase et al., 2012; Keitel et al., 2018; Kim et al., 2007), reflecting more stable phase relationship between the driving stimulus and SSVEPs when the flickering stimulus is attended. Taken together, removing the non-phase locked part of the signal and focusing on the phase-locked revealed a positive attentional modulation effect on alpha-band SSVEPs.

### Attention modulates second-harmonic SSVEP responses up to 110 Hz

Although the SSVEP waveform primarily contains the stimulus frequency (1*f*), harmonic components can also be present (2*f*, 3*f*, 4*f*, etc.), and are assumed to reflect nonlinearities in the response of the visual system (Labecki et al., 2016; Norcia et al., 2015). We investigated attentional modulation of responses at the 2*f* harmonic of the stimulus frequency in more detail, because 2*f* responses can also be modulated by attention, and sometimes even show stronger attention modulations as compared to the stimulus frequency (Kim et al., 2007; Kim et al., 2011; Vissers et al., 2017). Visual inspection of spectral response profiles revealed differences in the number and amplitude of harmonic responses across flicker frequencies (see Figure 3, middle panel). The number and magnitude of harmonics reflect deviations from sinusoidality of SSVEP response waveform, as purely sinusoidal SSVEP response would contain only one peak in the frequency spectrum. Therefore, as an exploratory analysis we quantified the attenuation of harmonic responses across flicker frequencies, with a special focus on comparison of harmonics between alpha (7-13 Hz) and non-alpha (3-6 Hz, 14-80 Hz) flicker frequencies.

The spectral response profile of the 2*f* harmonic contained only one peak at ~18 Hz, associated with responses to alpha-band flicker (Figure 7A). For the alpha-band flicker, the SNR of the 2*f* harmonic response was even stronger than that of 1*f* (see Figure 7C). For the gamma-band flicker, the peak in the 2*f* spectral response profile was absent (compare Figure 3A right panel and Figure 7A). This absence cannot be explained by the lower signal-to-noise ratio for higher flicker frequencies, as SNR of the 2*f* harmonic components was statistically higher than the null hypothesis for all flicker frequencies (p < .05, corrected for multiple comparisons across frequencies using cluster-based permutation testing).

**Figure 7.**
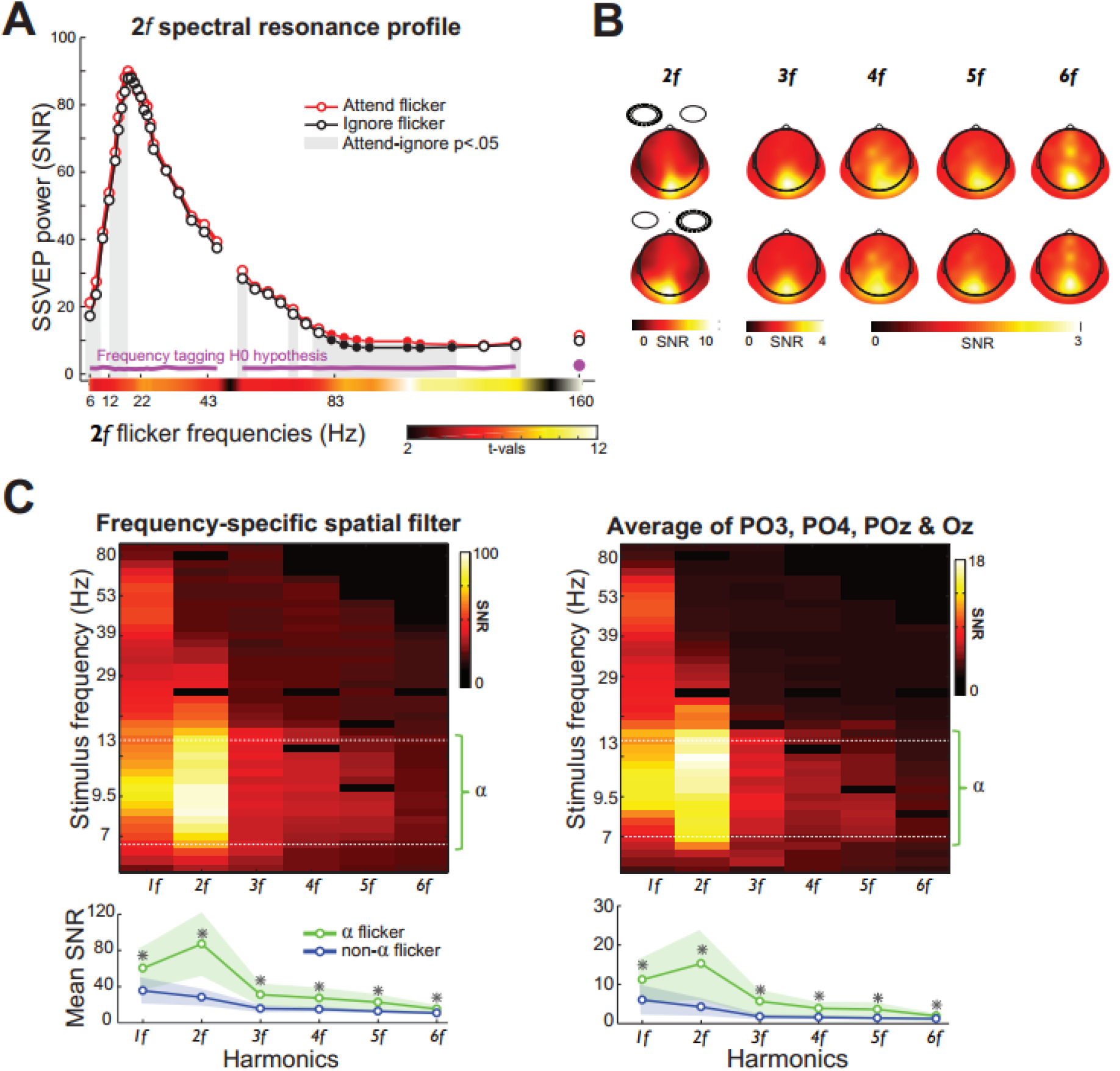
Higher harmonic components of SSVEPs. (A) Spectral profile of second harmonic (2*f*) component, plotted separately for “attend flicker” (red line) and “ignore flicker” (black line) conditions (collapsed across left-and right-hemifield flicker conditions). Conventions are the same as in Figure 3. (B) Subject-average RESS spatial filters of higher harmonic responses (2*f*, 3*f*, 4*f*, 5*f*, and 6*f*), constructed separately for each flicker frequency and separately for left- (top row) and right-flicker (bottom row) conditions. (C) SSVEP power at the stimulus and harmonically related frequencies, plotted separately for SSVEPs derived using frequency-specific spatial filters (left panel), and channel-level EEG time series. Line plots below represent average SNR at the stimulus frequency and harmonics for alpha- (8-13 Hz) and non-alpha flicker frequencies. Response magnitude for alpha-band flicker vs. non-alpha band flicker was statistically significant across all harmonic components (p < .001).

Statistically significant attentional modulation of 2*f* harmonic was constrained to low (up to 8.5 Hz flicker) and high flicker frequencies (35-70 Hz flicker), and numerically positive for all but 10 Hz and 10.5 Hz flicker frequencies (Figure 7A). Typically, robust attentional modulation of 2*f* responses is reported for pattern reversal stimuli (Kim et al., 2011), which by definition elicit only even harmonics of the stimulus frequency (Norcia et al., 2015). Thus, only a modest attentional modulation of the 2*f* harmonic might be due to the sine-wave stimulus luminance modulation we used.

We found differences in topographical maps of harmonic components (Figure 7B), such that topographies of higher harmonics were more anterior and localized as compared to more posterior and distributed topographies of lower harmonic responses (e.g. 2*f*; Figure 7B). The observed differences at the electrode level are consistent with previous reports on partial spatial segregation of harmonic SSVEP components at the source level (Fawcett et al., 2004; Heinrichs-Graham and Wilson, 2012; Pastor et al., 2007). Although 2*f* responses were present and significant for all flicker frequencies, 3*f* and higher harmonic responses were sharply attenuated for flicker frequencies above 30 Hz (see Figure 7C). Exploratory analyses revealed that alpha-band flicker elicited more and stronger higher harmonic components (alpha-band SSVEPs were significantly stronger as compared to non-alpha frequencies; Figure 7C, line graphs), meaning that alpha-band SSVEPs contained more non-sinusoidal features than non-alpha flicker SSVEPs.

### Alpha- and gamma-band flicker elicits maximally phase stable SSVEPs

To characterize fluctuations in the instantaneous phase of SSVEP response within each trial, we computed instantaneous SSVEP frequency – the first temporal derivative of single-trial phase time series (Cohen, 2014b). The observed fluctuations of instantaneous SSVEP frequency around the flicker frequency were more probable than could be expected in the absence of an external periodic drive (p < .05 for all flicker frequencies, corrected for multiple comparisons across frequencies). Importantly, flicker frequencies in the alpha and gamma bands that elicited the highest power SSVEPs were also associated with the smallest moment-to-moment variability in instantaneous SSVEP frequency, reflecting more stable phase of SSVEPs. Considering that instantaneous phase can be estimated more reliably when the power is high (Cohen, 2014b), the same resonance peaks in SSVEP power and SSVEP frequency fluctuation analyses may appear trivial (Figure 3 and Figure 8). However, more variable instantaneous SSVEP frequency for 47 Hz and 53 Hz flicker conditions, the estimates of which were influenced by high-power 50 Hz line noise, speak against this simple explanation (see Figure 8, square dots).

**Figure 8.**
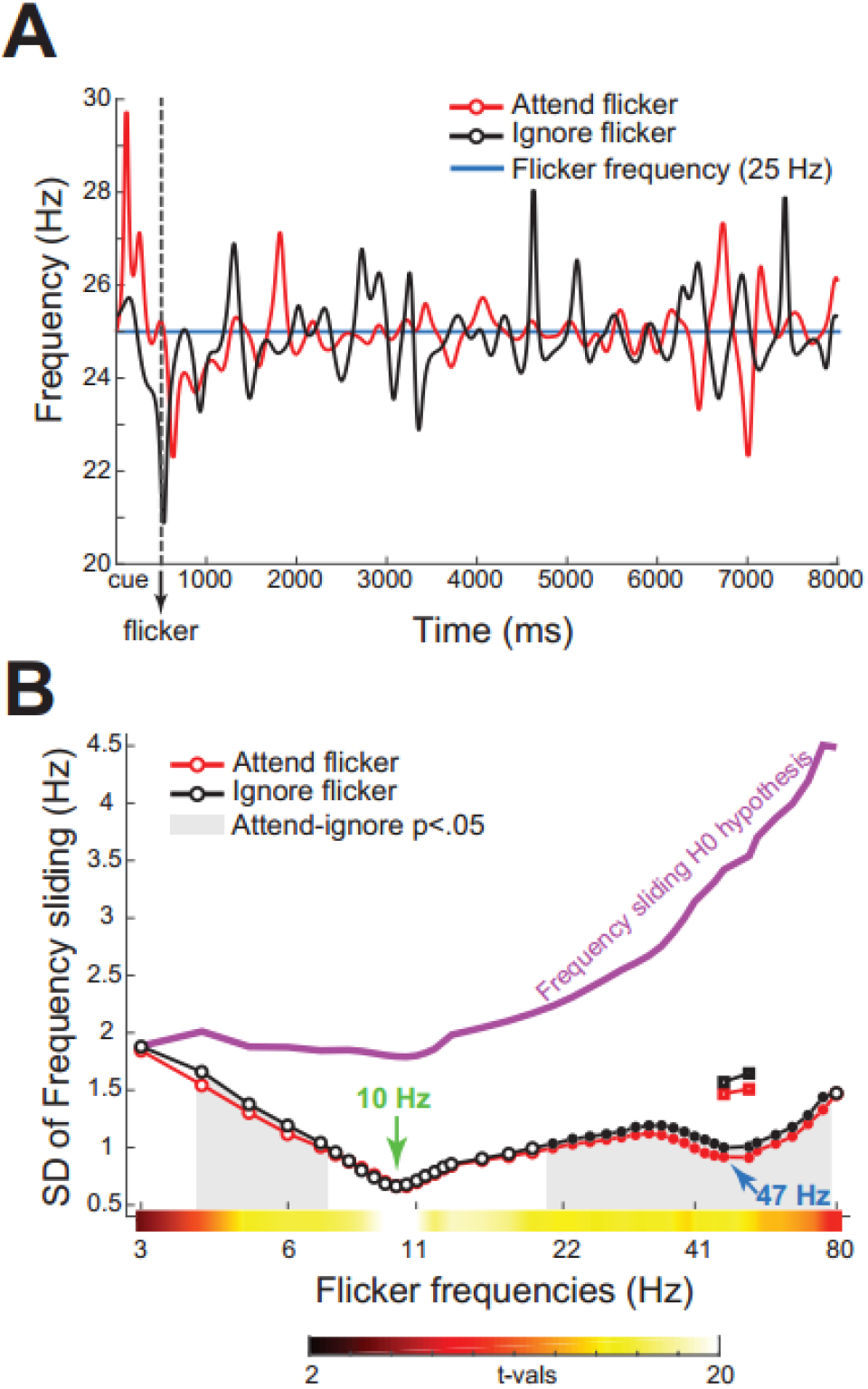
Fluctuations of instantaneous SSVEP frequency. (A) “Attend flicker” and “ignore flicker” exemplar trials illustrating non-stationarity of SSVEP response over time as reflected in fluctuations of instantaneous SSVEP frequency around the stimulus frequency (25 Hz). (B) Standard deviation (SD) of instantaneous SSVEP frequency changes over time as a function of flicker frequency. Conventions are the same as in Figure 3. Note that estimated variability of instantaneous SSVEP frequency for 47 Hz and 53 Hz flicker conditions was influenced by 50 Hz line noise (square dots), and thus was extrapolated from nearby flicker conditions (circle dots).

Attention decreased the variability of instantaneous SSVEP frequency for theta- and gamma-band, but not alpha-band flicker frequencies (see Figure 8). This may seem to contradict the phase-clustering analysis results, in which attention significantly reduced phase variability of alpha-band SSVEPs (Figure 6B). However, it is important to emphasize that phase-clustering analyses were based on average waveforms, whereas SSVEP frequency fluctuations were estimated on a single-trial level.

## DISCUSSION

We mapped out the effect of attention on SSVEPs generated in response to a wide range of temporal frequencies (3-80 Hz). The effect of attention on SSVEP amplitude was frequency-dependent: Positive for theta- (3-7 Hz) and gamma-band (30-80 Hz) flicker frequencies, but negative for alpha (8-13 Hz). Overall, SSVEP responses were significantly stronger and more phase stable at flicker frequencies in the alpha (7-13 Hz) and gamma (30-80 Hz) bands – the characteristic endogenous rhythms of the visual cortex. Covert spatial attention increased SSVEP amplitude to gamma band flicker on average by 20%, which is higher than attentional modulation of sustained gamma power in V1 in non-invasive MEG studies (5-10% in Magazzini and Singh, 2018), and comparable to invasive ECoG studies (~20% in Davidesco et al., 2013). Together, these findings have a series of theoretical and practical implications detailed below.

### Alpha- and gamma-band resonance of visual cortex to flicker stimulation

Corroborating previous qualitative reports on variability of SSVEP amplitude as a function of stimulation frequency (Bayram et al., 2011; Herrmann, 2001; Regan, 1968), we took a rigorous quantitative approach and we found statistically significant SSVEPs up to 80 Hz, with local maxima (resonance) in response to alpha- (~10 Hz, range 9-13 Hz) and gamma-band (~ 47 Hz, range 43-65 Hz) flicker. These resonance frequencies matched “natural” frequencies of the visual cortex: Population mean of resting-state alpha-band peak is centered at 10 Hz (Bazanova and Vernon, 2014), and peak frequency of visually induced gamma-band oscillations vary in the 42-65 Hz range (Muthukumaraswamy et al., 2010). Alpha and gamma resonance frequencies were uncorrelated across participants (Figure 4A), suggesting an independence of the two. While subject-specific resonance peaks in the alpha-band have been studied extensively (Fedotchev et al., 1990; Lazarev et al., 2001; Notbohm et al., 2016; Pigeau and Frame, 1992; Regan, 1989), to our knowledge this is the first time inter-subject variability in gamma-band resonance frequency is reported.

We found that not only SSVEP amplitude but also phase stability of SSVEPs is dependent on the temporal frequency of the visual input (Figure 6B and Figure 8B). Due to the deceivingly narrow-band representation of SSVEPs in the frequency domain, the frequency of SSVEPs is assumed to be stable over time (Norcia et al., 2015; Vialatte et al., 2010). Here, we show that the frequency of SSVEP, or SSVEP phase stability, fluctuates. We used two methods to evaluate SSVEP phase-stability: phase clustering (amplitude-independent measure of phase stability; Kim et al., 2007) and instantaneous frequency of SSVEPs (amplitude-dependent measure of phase stability; Cohen, 2014b). Despite the differences in mathematical implementation, the two methods essentially reflect how variable are the phase lags between the stimulus and the response. The two methods provided consistent results: stronger phase-locking to alpha-vs. non-alpha flicker frequencies (peak ~10 Hz, Figure 6B), and less variable instantaneous frequency of SSVEPs in response to alpha- and gamma-band flicker as compared to other frequencies (local peaks ~10 Hz and ~47 Hz in Figure 6B, and Figure 8B). Thus, not only amplitude but also phase-stability of SSVEPs is frequency-dependent.

Although phase synchronization with the stimulus may lead to power increase at the same frequency, this is not always the case (Schroeder and Lakatos, 2009). Increase in both phase coherence and power indicates either a superposition of stimulus-evoked and endogenous rhythmic activity (Lopour et al., 2013), or entrainment of endogenous oscillations by means of phase adjustment to the external periodic drive (Thut et al., 2011). Indeed, the entrainment and linear superposition of evoked responses are the two primary neural mechanisms put forward to explain the generation of steady state responses (for a recent review, see Zoefel et al., 2018). Irrespective of the debate concerning the neural mechanism underlying the generation of SSVEPs, the finding of individual gamma-band resonance frequency points to an exciting possibility to use flicker as a tool for studying interaction between exogenous and endogenous rhythms in the visual system at higher frequencies than has been reported before. However, additional research addressing the overlap between neural generators giving rise to gamma-band SSVEPs and visually induced gamma oscillations is needed (Hermes et al., 2017).

### Endogenous alpha may explain the reversed effect of attention on SSVEP power

Contrary to a commonly reported positive effect of attention on SSVEP amplitude (Andersen et al., 2011; Keitel et al., 2013; Morgan et al., 1996; Toffanin et al., 2009), we found that alpha-band flicker responses increased when attention was drawn away from the flickering stimulus. Increased SSVEP amplitude was accompanied by increased endogenous alpha power over the same hemisphere (i.e. contralaterally to the to-be-ignored flicker; Figure 5D). Co-occurrence of the two effects suggests a linear superposition of attention-related endogenous alpha oscillations and alpha-band SSVEPs at the electrode level, or an interaction between endogenous alpha oscillations and alpha-band flicker (entrainment). To dissociate alpha-band SSVEPs from endogenous alpha oscillations, we used a multivariate source separation approach and defined optimal spatial filters to isolate frequency-specific SSVEPs (Cohen and Gulbinaite, 2017). A prominent endogenous alpha peak was present in spatially filtered data regardless of the flicker frequency. The fact that endogenous alpha oscillations were partially rather than completely suppressed by SSVEP-specific spatial filters can be taken as evidence for a partial spatial overlap of the two EEG features.

We therefore leveraged temporal dynamics to better separate endogenous alpha from the flicker response, by isolating the phase-locked part of the SSVEP signal, thus removing the non-phase-locked endogenous alpha. This analysis approach revealed the expected direction of higher alpha-band SSVEP amplitude for attended vs. ignored stimuli. Attended flickering stimuli also showed more stable phase alignment of the SSVEP waveform to the driving stimulus, corroborating previous reports on increased inter-trial phase clustering of SSVEPs (Kashiwase et al., 2012; Keitel et al., 2017; Kim et al., 2007). Thus, attention had a positive effect on the phase-locked part of alpha-band SSVEPs. More generally, differences in attention effects on the phase-locked vs. total SSVEP signal indicates that the SSVEP signal is neither deterministic nor stationary (cf. Nunez and Srinivasan, 2006).

Under what circumstances could the negative attentional modulation of alpha-band SSVEPs be observed? The effect, to our knowledge, has been reported previously in two paradigms using central flicker stimuli (Chen et al., 2003; Ding et al., 2006; Wang et al., 2007). In both paradigms, broadly distributed attention evoked the expected pattern of higher SSVEP amplitude for attended stimuli, whereas narrowly focused attention evoked the reverse pattern of larger SSVEPs for ignored compared to attended stimuli. Thus, the paradoxical effect of attention on alpha-band SSVEPs was observed when interference from flicker and superimposed distractors was high. Based on these studies and our findings, we propose that reversal of alpha-band SSVEPs is due to active suppression of flicker, which is marked by an increase in endogenous alpha power (Jensen and Mazaheri, 2010; Klimesch et al., 2007; Mathewson et al., 2011). In support of this interpretation, some evidence suggests that alpha oscillations are more effectively entrained when alpha power is high (Kizuk and Mathewson, 2017). Thus, a prerequisite for observing negative attentional modulation of alpha-band SSVEPs is the presence of strong endogenous alpha oscillations to begin with, as well as high intensity of the entraining stimulus (Notbohm et al., 2016).

### Positive attentional modulation of gamma-band SSVEP amplitude and phase

Detection of gamma-band oscillations in non-invasive EEG and MEG studies highly depends on selection of optimal stimulus parameters (e.g. stimulus size, spatial frequency, contrast, eccentricity; Magazzini and Singh, 2018; van Pelt and Fries, 2013) and application of specific analyses techniques (Fries et al., 2008; Muthukumaraswamy and Singh, 2013), which can be limiting factors for studying gamma-band oscillations non-invasively. By applying optimal spatial filters to isolate frequency-specific SSVEPs (Cohen and Gulbinaite, 2017), we resolved robust gamma-band SSVEPs (up to 80 Hz, ~10 SNR units) and a subject-specific gamma resonance frequency (on average, 40 SNR) using a relatively small amount of data per flicker frequency (96 seconds).

Gamma-band SSVEPs were positively modulated by attention. The topographical peak of attentional modulation was centered over occipitoparietal areas contralateral to the flickering stimulus (Figure 3B). Although multiple harmonic SSVEP components precluded us from analyzing lateralization of endogenous gamma-band oscillations, gamma power generally is increased contralaterally to the direction of attention (Doesburg et al., 2008; Fries et al., 2001; Green et al., 2017; Magazzini and Singh, 2018). Thus, there is a topographical overlap between gamma-band SSVEPs and attention-related endogenous gamma. Phase-based analysis of instantaneous SSVEP frequency also revealed a positive effect of attention (Figure 8B), such that phase-alignment to the gamma-band driving stimulus was stronger for attended stimuli. Finding a subject-specific gamma resonance frequency and an attentional modulation of gamma-band SSVEPs opens up new possibilities for tracking attention with high temporal precision and testing the causal effects of flicker-driven gamma-band responses on perception and attention non-invasively.

### Frequency-specific flicker effects on cognition and perception

Although it is implicitly assumed that rhythmic visual stimulation does not have frequency-specific cognitive effects (Keitel et al., 2014), some studies provide evidence for behavioral consequences of alpha-band flicker (de Graaf et al., 2013; Gulbinaite et al., 2017; Williams, 2001). Here, we observed frequency-specific cognitive and perceptual effects of flicker, with non-overlapping frequencies affecting the two processes. On a cognitive level, targets were more accurately detected when the to-be-ignored hemifield flickered in low (theta and alpha) or high (gamma) frequencies. On a perceptual level, target stimuli needed to be brighter to be detected when the background flickered at ~15 Hz. However, target detection was improved when the unattended hemifield flickered at ~30 Hz. These results can be partially explained by frequency-dependent enhancement of perceived stimulus brightness: For sinusoidal flicker, as the one we used, brightness enhancement peaks at ~16 Hz (Wu et al., 1996), for square-wave flicker the peak is at ~8 Hz (Bartley, 1938; but see Bertrand et al., 2018). Given these non-linearities in brightness perception of flicker, blue targets presented on ~15 Hz flickering background seemed dimmer due to increase in perceived brightness of the background. Relatedly, static stimuli appear dimmer than flickering stimuli of the same mean luminance (“Brücke-Bartley Effect”, Bartley, 1938). Considering that the luminance of the static background was kept constant, it is possible that some flicker frequencies made the static background appear relatively dimmer, thus affecting the contrast between blue targets and white background. Taken together, these results indicate frequency-specific cognitive and perceptual effects of flicker even when it is present in remote parts of the visual field.

### Harmonic components of SSVEPs – a method to study ultra-high frequency activity with EEG

High-frequency tagging allows to measure temporal dynamics of cognitive processes on a finer temporal scale, and is arguably less tedious for the participant. However, due to low signal-to-noise ratio (Herrmann, 2001), high-frequency tagging has rarely been explored. Here, we resolved the SSVEP signal with >10 SNR for frequencies up to 160 Hz (second harmonic component of 80 Hz flicker), thus providing a robust method for studying ultra-high frequency activity in EEG.

Alpha-band flicker elicited the strongest harmonic responses and response amplitudes at the 2*f* component (i.e.18 Hz) were ~10% higher than that of the stimulus frequency (i.e. 9 Hz). The prominent alpha-band flicker harmonics can be explained by at least two non-mutually exclusive mechanisms. First, non-sinusoidal features in alpha-band SSVEPs could reflect waveform non-sinusoidalities of endogenous alpha oscillations (Cole and Voytek, 2017; Lozano-Soldevilla et al., 2016; Mazaheri and Jensen, 2010; Schaworonkow and Nikulin, 2018). This also suggests that more effective entrainment of endogenous alpha-band oscillations could be achieved using non-sinusoidal stimulation protocols (Hutt et al., 2018). Second, already at the level of the retina responses to alpha-band flicker contain a stronger 2*f* harmonic component as compared to other frequencies (Burns et al., 1992; Odom et al., 1992).

### Conclusions

The range of frequencies used for “frequency-tagging” stimuli in the studies of selective attention varies considerably: From 3 Hz to ~30 Hz. Does attention modulate SSVEP amplitude at higher flicker frequencies and are all tagging frequencies equally “useful” for tagging selective attention? We found that attentional modulation of SSVEP amplitude is the most pronounced in alpha and gamma (30-80 Hz) frequency bands at subject-specific resonance flicker frequencies, with respectively negative and positive effects of attention. Flicker also had frequency-specific cognitive and perceptual effects. Together, these findings speak against frequency neutrality when using flicker to study selective attention, and demonstrate that some tagging frequencies are more useful than others.

## ACKNOWLEDGEMENTS

This research was funded by the ERC-CoG P-CYCLES (N°614244) awarded to Rufin VanRullen. We thank Ludovic Tyack for his assistance in designing and building the LED stimulation device, and Dr. Kenneth Knoblauch for measuring luminance of the stimuli.

## SUPPLEMENTARY MATERIALS

**Figure S1.**
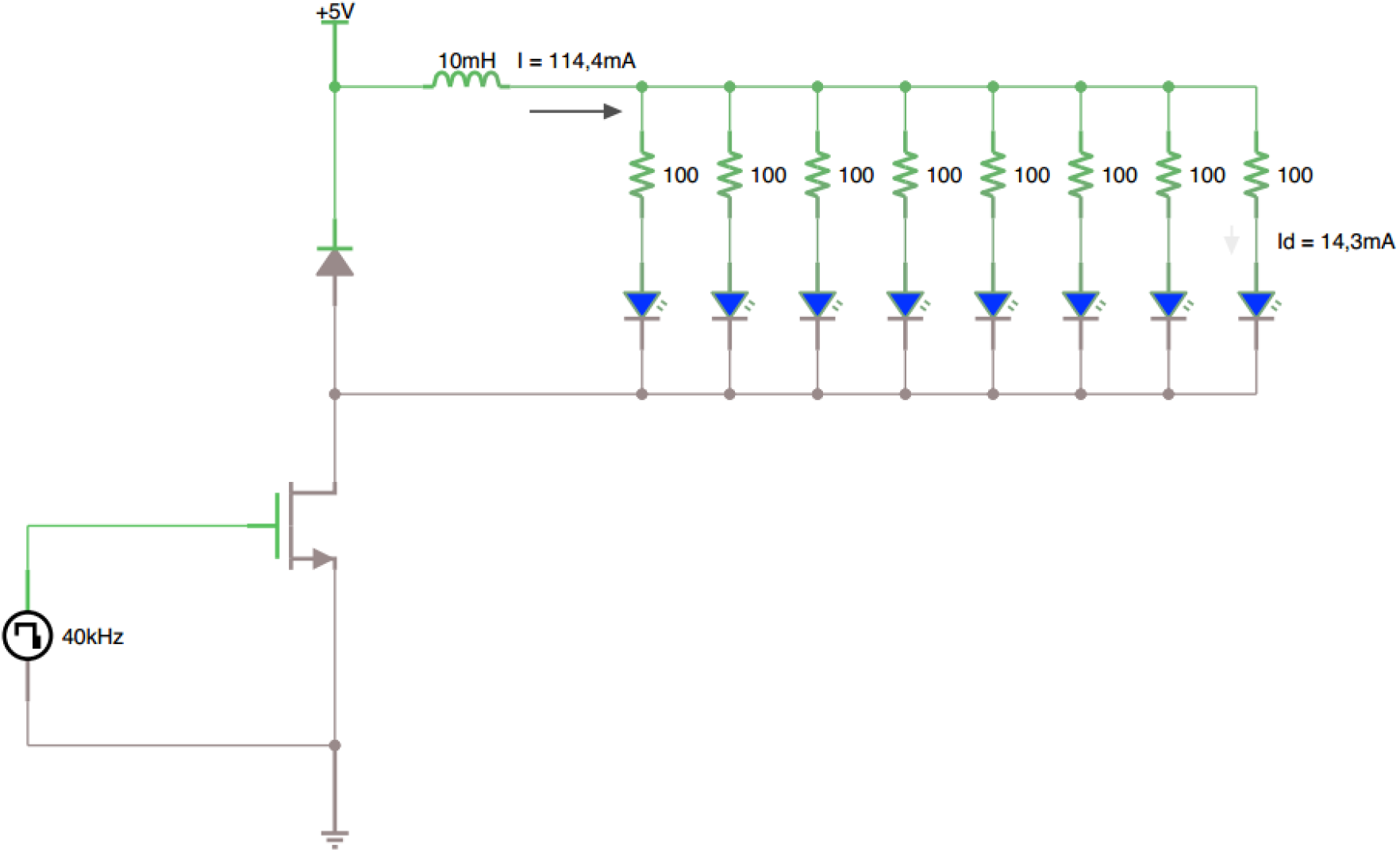
A circuit diagram of the sinewave generator.

